# Soft extracellular matrix drives an endoplasmic reticulum stress-dependent S quiescence underlying molecular traits of pulmonary basal cells

**DOI:** 10.1101/2023.08.18.552093

**Authors:** Pierre-Alexandre Laval, Marie Piecyk, Pauli Le Guen, Mirela-Diana Ilie, Joelle Fauvre, Isabelle Coste, Toufic Renno, Nicolas Aznar, Celine Hadji, Camille Migdal, Cedric Duret, Philippe Bertolino, Carole Ferraro-Peyret, Alice Nicolas, Cedric Chaveroux

## Abstract

Cell culture on soft matrix, either in 2D and 3D, preserves the characteristics of progenitors. However, the mechanism by which the mechanical microenvironment determines progenitor phenotype, and its relevance to human biology, remains poorly described. Here we designed multi-well hydrogel plates with a high degree of physico-chemical uniformity to reliably address the molecular mechanism underlying cell state modification driven by physiological stiffness. Cell cycle, differentiation and metabolic activity could be studied in parallel assays, showing that the soft environment promotes an atypical S-phase quiescence and prevents cell drift, while preserving the differentiation capacities of human bronchoepithelial cells. These softness-sensitive responses are driven by defects in proteostasis and enhanced basal endoplasmic reticulum stress. The analysis of available single cell data of the human lung also showed that this non-conventional state coming from the soft extracellular environment is indeed consistent with molecular feature of pulmonary basal cells. Overall, this study demonstrates that mechanical mimicry in 2D culture supports allows to maintain progenitor cells in a state of high physiological relevance for characterizing novel molecular events that govern progenitor biology in human tissues.

## 1. Introduction

As cells sense extracellular mechanics, interest in extracellular forces on cell behavior and fate has increased significantly in the recent years. Mechanical signals from soft environment in 2D and 3D directly influence cell state by favoring stemness features, and, subsequently, their response to extracellular cues including therapeutic and microenvironmental stresses [1,2]. By extension, the relevance of regular tissue culture plastic (TCP) has been questioned as this matter displays an extraphysiological rigidity (in the gigapascal range) that promotes abnormal cell polarization and loss of differentiation potential [3,4]. Furthermore, prolonged culture on TCP is associated with cell drift along passages, a well-known issue underlying the reproducibility crisis [5]. Similar to 3D culture, hydrogel-coated plates represent a biocompatible support capable of reproducing the mechanical properties of the tissue of interest while maintaining the progenitor state [6]. These 2D supports, whose compliance is close to that encountered by cells *in vivo,* are similar in use to regular TCP and limit the heterogeneity in cell assemblies often encountered in 3D [7]. The possibility of amplifying cells at a large scale and its compatibility with commercially available live cell imagers confers a decisive asset for using this approach in mechanistical studies.

However, the fabrication of plates with soft bottoms is far from standard and although efforts have been put in the last 15 years to design soft bottom multi-well plates, critical issues remain. Two main difficulties were identified: i) the need to have a reduced variability in the mechanical properties and surface chemistry between wells, while ii) ensuring no fluid leakage and exchange between wells. Several developments have attempted to address these issues, including polymerization of the hydrogel well per well or fabrication of hydrogel spots to be enclosed by a multiwell frame [8],[9]. While the first strategy makes it difficult to control: i) the flatness, ii) the interwell reproducibility, iii) the absence of unpolymerized toxic residues and iv) the thickness of the hydrogel layer[9] the second strategy meets the difficulty of sealing the engineered bottom to the upper frame. For the latter, controlling the absence of interwell leaks is critical and the proposed solution of clamping the both parts at the edges of the plates does not provide robust mitigation owing to the deformability of the upper frame [10].

These technological issues contribute to the overall poorly addressed question of whether tissue culture on soft supports and associated cellular reprogramming truly reflect the cell biology in human. While there is extensive literature documenting the differences between cellular responses on soft substrates and TCP (for review [11]), it remains to be assessed whether cells grown on mechanomimetic supports not only behave differently from those grown on TCP but also better retain the physiological characteristics. In terms of phenotypic impact, a large number of studies have reported that culture on biomimetic hydrogels represses cell proliferation and favors a quiescent phenotype associated with the maintenance of pluripotency in numerous models compared to standard culture conditions at early passages [12,13]. Pluripotent epithelial cells possess the ability to differentiate into a specific cell type upon specific cues. Consistently, in adult tissue, the number of proliferative cells is rare, indicating that the majority of cells reside in a quiescent or senescent state to preserve their functional features [14]. Loss of quiescence leads to stem cell depletion and defects in tissue regenerative capacity illustrating the tight connection between cell identity and the proliferative state. The term “quiescence” is widely used to designate a reversible non-proliferative state enabling cells to re-enter the cell cycle [15]. Although often restricted to exiting the G1 phase towards G0, numerous types of quiescent states have been described along the cell cycle [16–18]. Few reports suggest that mechanical cues halt cells but not in the G0 phase, suggesting that an atypical quiescence might underlie progenitor states in cells grown on hydrogel-based supports [19,20]. Furthermore, the fundamental mechanisms by which soft supports trigger quiescence is still unknown. Cell quiescence is classically triggered following microenvironmental cues, including nutritional deprivation. This functional analogy between cells subjected to metabolic stress or grown on soft matrix suggests that common pathway(s) related to nutrient sensors might be involved in the mechanically driven modification of cell fate. The endoplasmic reticulum (ER) is considered as a nutrient-sensing organelle [21]. Upon nutritional stresses, the subsequent accumulation of misfolded proteins within the ER lumen leads to the ER stress (ERS) and activation of the relative signaling pathways called unfolded protein response (UPR). The UPR then orchestrates of a complex transcriptional, post-transcriptional and translational reprogramming that contributes to the restoration of cellular homeostasis and plasticity. Nonetheless, the ERS molecular axes are now emerging as key molecular events also implicated in the sensing of mechanical cues and cell identity changes [22]. Whether cells grown on soft matrices display proteostasis defects, dysregulated ERS signaling that might support cell cycle arrest and phenotypic modifications remains unknown.

Therefore, in the present study, we introduce a novel technology for the generation of multiwell hydrogel plates that drastically improves the interwell homogeneity and reliability of the desired stiffness compared to existing methods. Based on this innovation, we aimed to: i) define whether slow proliferative capacities on mechanomimetic environment are associated with the acquisition of a specific type of quiescence supporting the progenitor state, ii) determine whether these soft supports enable long-lasting cell culture with reduced cell drift, and iii) define whether a chronic ER stress controls this progenitor phenotype, iv) validate in human tissue the relevance of the findings. Using human bronchoepithelial cells (HBEC-3KT) as a model of pulmonary cells and the soft multiwell plates to conduct imaging, proteomic and transcriptomic assays in parallel on the same cellular batch, we show that cell culture on a mechanomimetic matrix with stiffness comparable to lung tissue favors a quiescent state in the S phase preserving differentiation capabilities along passages. We also demonstrate that Ca^2+^ depletion in the ER, subsequent chronic proteostasic and phosphorylation of eIF2a drive this non-canonical quiescence state. Our interdisciplinary study thus shows for the first time that stemness maintenance by mechano-mimetic environment is driven by the UPR. Furthermore, by analyzing single cell databases, we show that this observation is consistent with the biology of pulmonary basal cells in the human lung and resistance to ERS-mediated cell death caused by metabolic stress.

## 2. Results

### 2.1. Development of a new methodology for improving stiffness reliability in multiwell hydrogel plates

Reliable and comparable evaluation of the biological substratum governing cellular dynamics within hydrogels hosted in multiwell plates mandates uniform stiffness prevailing across distinct well compartments within a given plate. As described above, the current limitations implicated the development of a new strategy to ensure softness homogeneity across the different wells of the plate. Inspired from the work of Ahmed et al, engineering soft supports in the 6-well plate format was prioritized. This format not only serves as a convenient platform for real-time imaging but also furnishes ample biological matter to facilitate subsequent biochemical and molecular analyses [10]. First a polyacrylamide hydrogel layer was uniformly UV-cured on a multiwell-sized glass base treated to allow the covalent binding of the hydrogel. Before UV-exposure, the prepolymer solution was covered by a transparent hydrophobic glass slide equipped with 40 µm high wedges (Fig. 1A). Thus, the polymerized hydrogel had a flat surface and exhibited a constant thickness throughout. Tuning the exposure time allowed to tune the stiffness of the hydrogel. Extensive rinsing could be performed to remove unreacted compounds [10]. In contrast to the clamping method proposed by Ahmed et al.[10], the hydrogel layer was sealed to the bottomless multi-well frame using a non-toxic liquid adhesive (Fig. 1B). The glue was penetrating in the depth of the hydrogel thus guaranteeing a water-tight barrier in between the wells. To address the impact of soft matrix on pulmonary cell biology, 6-well plates were generated with 3 kPa polyacrylamide bottoms whose stiffness is in the range of physiological pulmonary rigidities measured in human samples of the order of 3 kPa [23,24]. Atomic force microscopy measurements confirmed that this technology enabled a high degree of uniformity within every well and in between the wells, with a Young’s modulus of 3.3+/-0.5 kPa (noted 3 kPa hereafter for the sake of simplicity) (Fig. 1C). Furthermore, as polyacrylamide is not permissive to cell adhesion, the surface of the hydrogel was then coated well per well with collagen I (1 µg/cm^2^), a major component of the lung matrisome [25]. The uniformity of matrix coating was achieved by evaporating the protein solution as described by Palva et al [26] until the hydrogel surface was dried and confirmed by immunofluorescence (Fig. 1D). TCP wells were also coated with the same matrix for establishing the phenotypic contribution of the substrate stiffness only, without introducing any bias through the biochemical environment. Since the starting material is a large hydrogel plate instead of polymerizing the hydrogel well per well, this novel approach can be extended to smaller multiwell soft supports and offers the possibility to fabricate large amounts of plates at once without apparent interwell leakage (Supplementary fig. 1A, B).

**Fig. 1.**
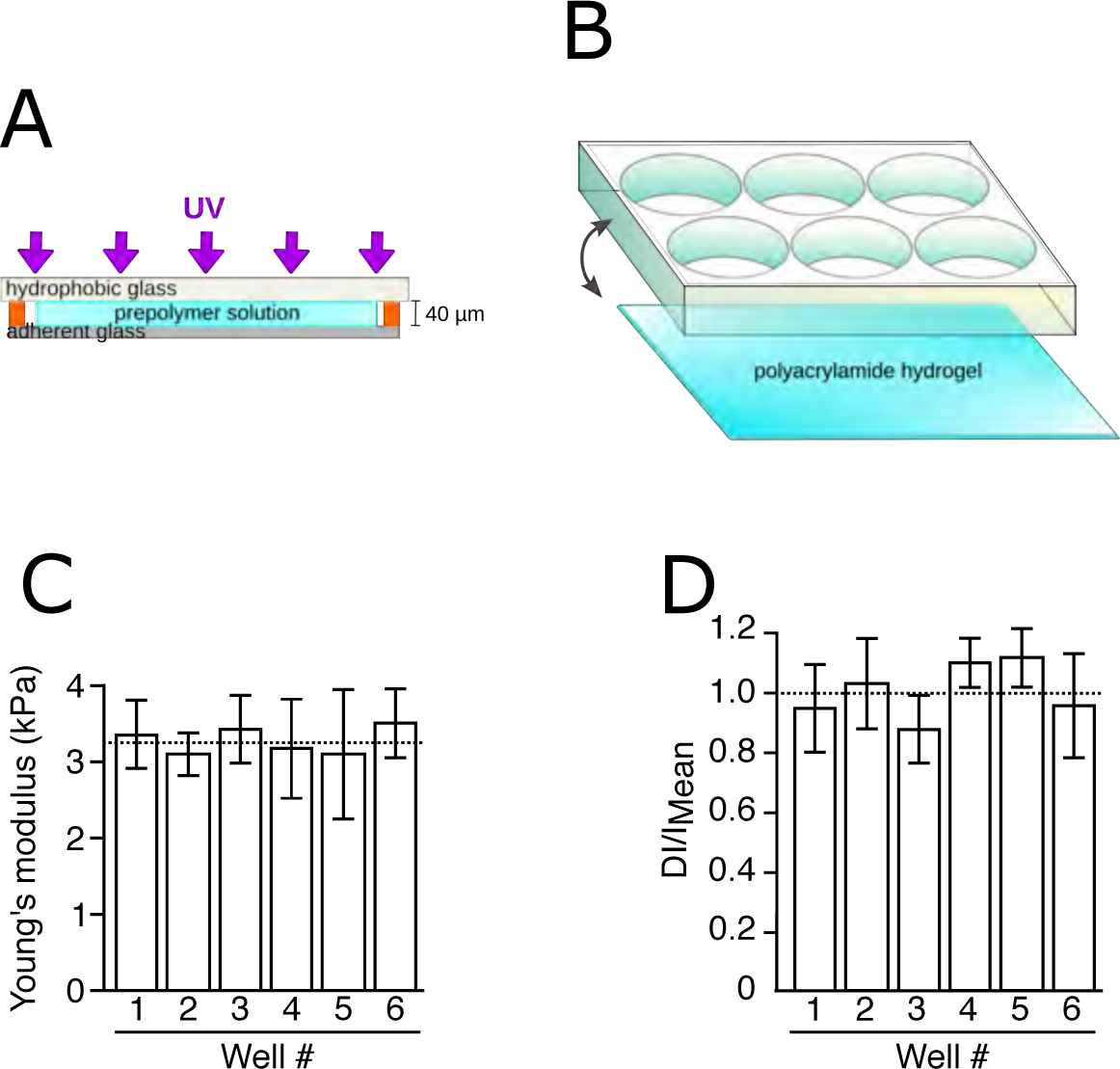
Design of multiwell plates with homogenous pulmonary stiffness across wells. A. A 40 µm thick layer of polyacrylamide prepolymer solution is UV-cured on a large glass slide. B. The hydrogel coated glass is further bonded to a bottom-less plate with a liquid adhesive that penetrates in the depth of the hydrogel and makes a physical barrier between the wells. C. Stiffness characterization of the soft bottom in each well. Young’s modulus: 3.3 +/-0.5 kPa. D. Relative variation of the surface density of the collagen 1 that is grafted onto the hydrogel in each well. Variation is of +/-0.13.

### 2.2. Pulmonary mechanomimetic supports promote an atypical quiescence in S phase

To investigate the impact of mechanomimetic hydrogels on lung cell identity at early and late passages, human broncho-epithelial cells (HBEC-3KT) were seeded on regular TCP dishes or on the flat 3 kPa polyacrylamide hydrogels (Fig. 2A). Confluency measurement confirmed the compatibility of our innovation with classical live-imager similarly to TCP supports and revealed that cells grown on soft supports were proliferating much slower than those evolving on TCP (Fig. 2B, C). We found that less than 10% of cells were stained for the proliferation marker KI67 on soft supports compared to 60% on TCP (Fig. 2D). The number of KI67-positive cells found on 3 kPa were closely similar to the number of KI67-positive epithelial cells as previously described in the human normal lung (11.3 %) suggesting that cell culture on a 3 kPa soft substrate better mimics the pulmonary physiology [27]. In addition, cells remained active since the mobility was not affected by the soft hydrogel compared to regular plastic support (Fig. 2E). To determine whether this slow-proliferating phenotype is maintained in long-term culture, HBEC-3KT were maintained either on TCP or 3 kPa supports for 20 passages and the doubling time was determined every 5 passages (Fig. 2F). Although the doubling time of these cells was about 2 days on plastic, it drastically increased up to about 7 days once the cells were maintained on 3 kPa supports. In both cases, this index remained fairly stable over time, regardless the stiffness of the supports. Slow/non-proliferating cells are often qualified as quiescent or senescent depending on their capabilities to progress again in the cell cycle once put back in proliferative conditions. Non-proliferating cells phenotypically often display cellular enlargement [28]. Consistently, HBEC-3KT cells grown on 3 kPa showed a significant increase of cell area and this phenotype was conserved even after 20 cell passages (Fig. 3A). Gene expression analysis of common canonical markers of quiescence and senescence (*P16*, *P21*, *P27* and *FOXO3A*) confirmed the change of cell state (Fig. 3B). To determine whether these changes are associated to a higher degree of senescence, a ß-galactosidase assay was performed (Fig.3C). We observed that cell culture on the soft matrix increased significantly the number of senescent cells compared to TCP. Nevertheless, this fraction represented only 12% of the total population indicating that, on soft substrate, the majority of HBEC-3KT cells were blocked in a quiescent state. The precise localization of quiescent HBEC-3KT within the cell cycle was analyzed by flow cytometry. Figure 3D shows that HBEC-3KT grown on soft supports halted in S phase whereas we observed a drastic drop of cells localized in the G0/G1 phase compared to the TCP condition. However, no difference was found between the two conditions regarding the proportion of cells in G2/M phase. These experiments illustrate that HBEC-3KT grown on soft supports display an atypical quiescent state in S phase instead of the canonical G0 arrest. To assess whether these cells maintain their capacity to re-enter into a proliferative state, the cells were grown one passage on TCP or 3 kPa supports, then cross-seeded either on 3 kPa or on TCP supports and the confluency index was measured (Fig. 3E). As expected, a severe repression of proliferation was observed when the cells were grown on 3 kPa supports independently of the support of origin. However, when cells grown on soft hydrogel were transferred to TCP, they regained in large part their proliferative capacity.

**Fig. 2.**
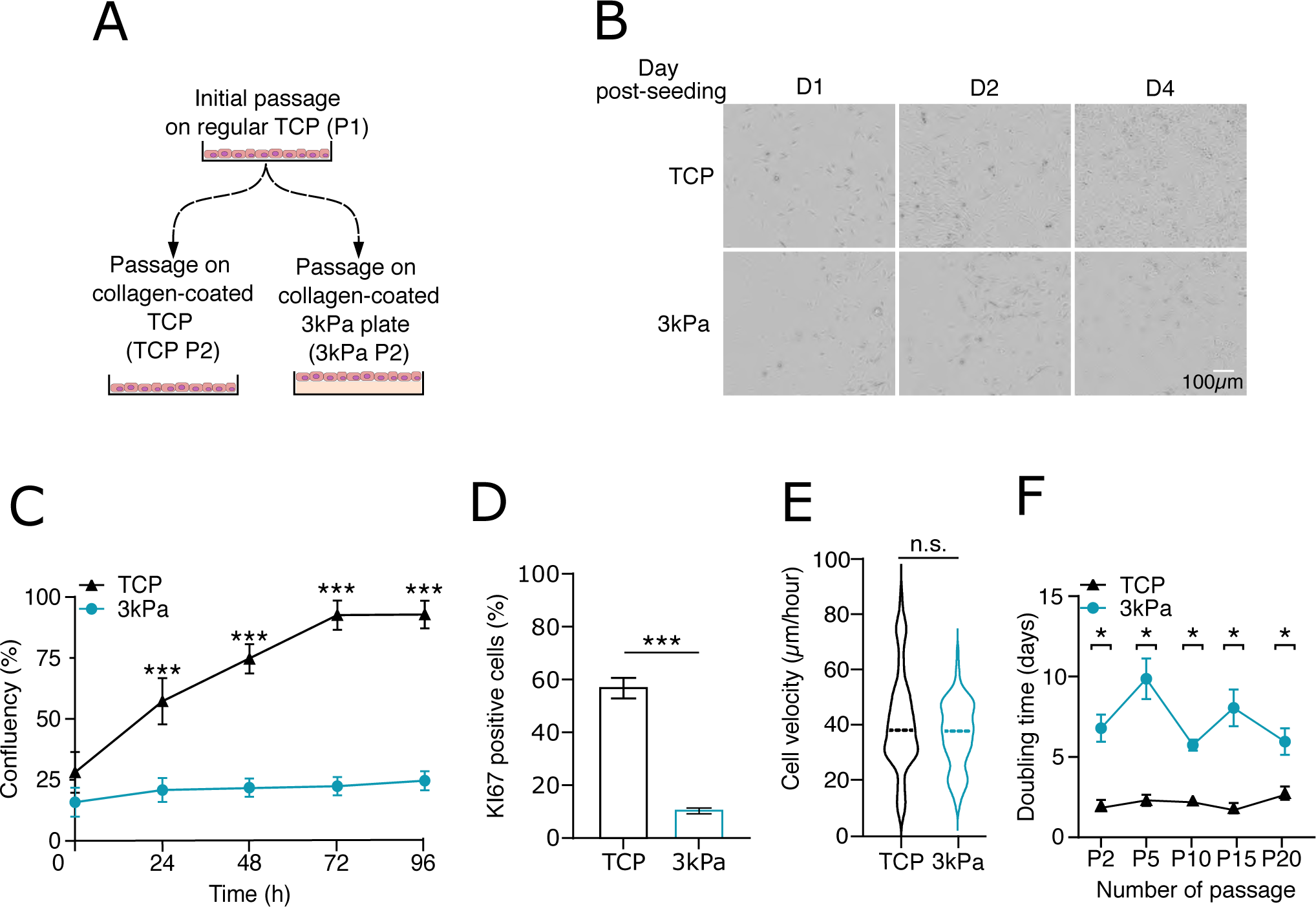
Soft hydrogels hinder human bronchoepithelial cells proliferation. A. Schematic representation of serial passages done on either Tissue Culture Plastic (TCP) or 3kPa hydro-gel. B. Pictures from HBEC-3KT cells grown on either TCP or 3kPa stiffnesses and followed over 4 days. C. Cell confluency measurement of HBEC-3KT cells grown on either TCP or 3kPa stiffnesses confluency over 4 days of culture. Data are expressed as mean +/-s.e.m of independent experiments (n = 3), Two-way ANOVA and Tukey’s multiple comparisons test were performed (***p < 0.001). D. Quantification of the proliferative subpopulation (Ki67-positive cells) of HBEC-3KT cells cultured on either TCP or 3kPa stiffnesses for 2 days. Data are expressed as mean +/-s.e.m of independent experiments (n = 3), Student’s t-test, *** p < 0.001. E. Cell velocity measurement of HBEC-3KT cells cultured on either TCP or 3kPa stiffnesses for 2 hours. Distribution is shown as violin plot. Data are expressed as mean +/-s.e.m of independent experiments (n = 3), Student’s t-test. F. Doubling time measurement of HBEC-3KT cells grown on either TCP or 3kPa stiffnesses over 20 pas-sages. Data are expressed as mean +/-s.e.m of independent experiments (n = 3), Student’s t-test, * p < 0.05.

**Fig. 3.**
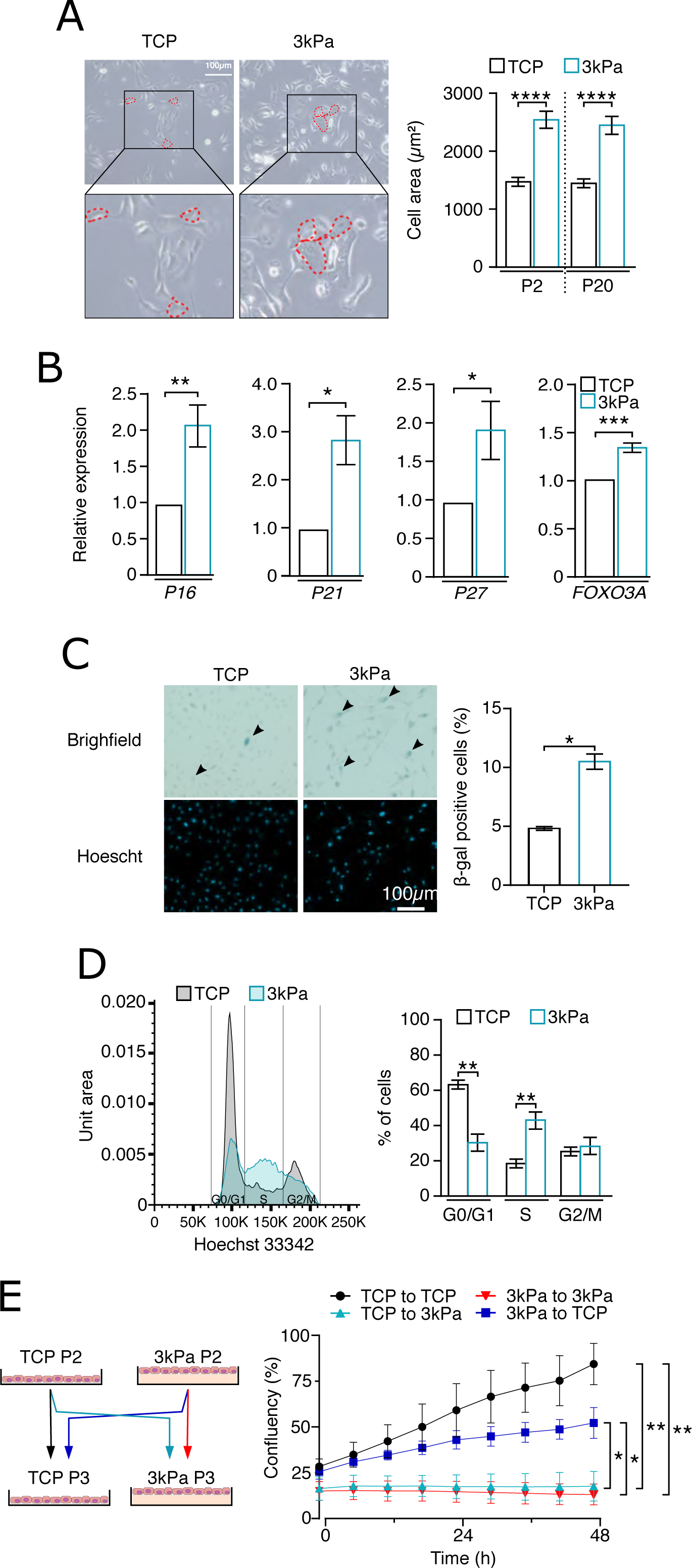
Pulmonary soft hydrogel induces a reversible quiescence stage in S-phase. A. Cell area measurement of HBEC-3KT cells grown on TCP or 3kPa stiffnesses. After 2 days of culture on both supports, cell edges were delimitated (outline in red, right panel) and cellular area were measured (left panel). Data are expressed as mean +/-s.e.m of independent experiments (n = 3). Student’s t-test (*p < 0.05) B. Gene expression analysis of canonical markers of quiescence and senescence *P16*, *P21, P27* and *FOXO3A* in cells grown on either TCP or 3kPa stiffnesses. Data are expressed as mean +/-s.e.m of independent experiments (n = 4), unpaired two-tailed t test (*p<0.05; **p< 0.01; ***p<0.001). C. Senescence measurement of HBEC-3KT cells cultured on TCP or 3kPa stiffnesses. After 2 days of culture, SA-β-galactosidase assay was performed, and cells were imaged (left panel). The percentages of positive cells were quantified (right panel). Data are expressed as mean +/-s.e.m of independent experiments (n = 3), Student’s t-test, (*p<0.05) D. Cell cycle analysis by flow cytometry of cells cultured on TCP or 3kPa stiffnesses. Cell distribution was plotted according to DNA content (left panel) and quantification of each cell cycle groups was assessed (right panel). Data are expressed as mean +/-s.e.m of independent experiments (n = 3), paired two-tailed t test (**p< 0.01) E. Quiescence reversibility assay of HBEC-3KT cells according to the described conditions. Cells were seeded on either TCP or 3kPa stiffnesses and then cross-platted following the schematic (left panel). Confluency was then assessed over 48 hours using live-imaging system. Data are expressed as mean +/-s.e.m of independent experiments (n = 3). Two-way ANOVA and Tukey’s multiple comparisons test were performed. Only the final time point statistics is presented (*p<0.05; **p< 0.01)0

### 2.3. Human bronchoepithelial cells grown on the soft supports show limited phenotypical drift after 20 passages

Similarly to our *in-vitro* observations, progenitor cells reside in a quiescent state in the adult normal tissue [29,30]. Thus, we sought to determine whether growing HBEC-3KT on soft matrix preserves their capacity to differentiate. This cell line has the capability to form alveoli once the cells are transferred on air-liquid interface (ALI) system. Thus, we tested whether the support rigidity and its associated cell identity changes the HBEC-3KT capacity of alveogenesis (Fig. 4A). Interestingly, when the cells are grown on an ALI membrane, which is a stiff porous membrane, the increased cell area caused by the soft support is maintained regardless of the number of passages, indicating that growing cells on the rigid ALI membrane does not reverse this phenotype (Fig. 4B). However, we noticed that, for both conditions, cell area at late passage (P20) was lower than at the early passage (P2), which could suggest that cells underwent a phenotypic drift in both culture conditions. At P20, soft supports still exhibited significantly larger cells than the TCP counterparts. We then evaluated their phenotype drift by analyzing the evolution of their differentiation potential along passages. This was first done by measuring the expression of classical markers of pulmonary progenitors (*KRT5*, *KRT14* and *TP63*) (Fig. 4C). At P2, the expression level of these genes was not significantly different between the two types of supports. However, at P20, this was significantly reduced for cells grown on TCP whereas it was maintained in cells grown on 3 kPa supports. This indicates that cell culture on 3 kPa, but not on TCP, maintains the HBEC-3KT pool of progenitors and differentiation potential. Then, a differentiation protocol was conducted. After 21 days, the ALI membranes were harvested and histological assessment of alveoli formation was performed (Fig. 4D). Consistent with the gene expression assays, we did not observe a significant difference between the two culture conditions in the ability of HBEC-3KT cells to generate alveoli at P2. However, at P20, the cells beforehand grown on TCP failed to generate alveoli whereas this ability was maintained for the cells previously grown on 3 kPa. At the transcriptional level, although the expression ratio of the alveolar markers (*PDPN* and *ACE2*) was not different in the differentiated tissues from P2, this was strongly increased in differentiated alveolar tissue from cells grown on soft compared to TCP at P20 (Fig. 4E)

Overall these data demonstrate that mechanomimetic substrates promotes the acquisition of a quiescent phenotype in S phase. This non-canonical type of quiescence is associated to the preservation of progenitor pool and differentiation capabilities of pulmonary cells along passages.

**Fig. 4.**
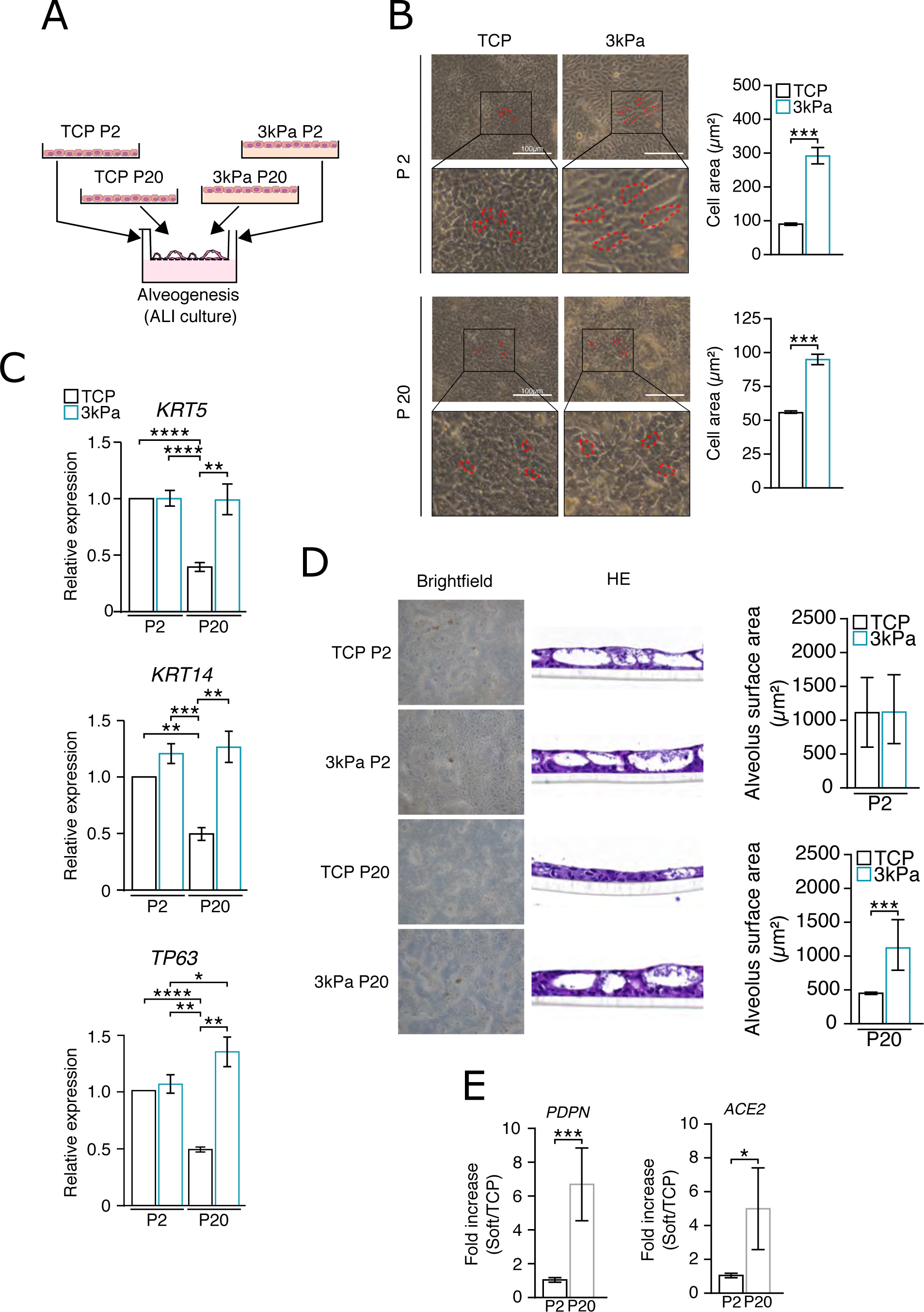
Pulmonary soft hydrogel retains cell differentiation capabilities over high number of passages. A. Schematic representation of Air-Liquide Interface (ALI) culture of HBEC-3KT Cells cultured over short (2 passages) or long (20 passages) periods of time on Tissue Culture Plastic (TCP) or 3kPa plates. B. HBEC-3KT cells were imaged before air-lift (induction of differentiation) for each condition (left panel) and cell area was measured (right panel). Data are expressed as mean +/-s.e.m of independent experiments (n = 3), student t test (***p<0.001) C. RT-qPCR analysis of progenitor markers *KRT5*, *KRT14* and *TP63* in HBEC-3KT cells pre-cultured on either TCP or 3kPa plates and seeded in ALI membranes (before airlift). Data are expressed as mean +/-s.e.m of independent experiments (n = 3), one-way ANOVA and Tukey’s multiple comparisons test (**p< 0.01; ***p<0.001). D. ALI cultures of HBEC-3KT cells were imaged after 21 days of differentiation and stained with Hematoxylin-Eosin (HE), highlighting alveolar-like structures. Quantification of the alveoli area are indicated on the right panel. Data are expressed as mean +/-s.e.m of independent experiments (n = 3), student t test (***p<0.001). E. RT-qPCR analysis of alveolar cell markers *PDPN* and *ACE2* in HBEC-3KT ALI culture 14 days post air-lift. Relative fold increase was assessed based on passage number. Data are expressed as mean +/-s.e.m of independent experiments (n = 3), student t test (*p< 0.05; ***p<0.001).

### 2.4. Cells grown on soft supports display alteration of protein synthesis but not DNA damages

Then we sought to identify the mechanisms limiting the S/G2 transition. Although several molecular and metabolic pathways are implicated in the maintenance of a G0 quiescent state, those supporting this atypical state of quiescence are poorly described. Activation of the DNA damage response (DDR) and particularly the ATR signaling pathway constitute a cell cycle checkpoint regulating the S/G2 transition [31]. The ATR signaling pathway is classically activated by DNA replication stress leading to the phosphorylation of Chk1. However the ATM-KAP1 axis has been suggested to also being activated upon replication stress induced by a metabolic stress [32]. To determine whether induction of the DDR pathways impairs the S/G2 transition, canonical markers of both axes have been assessed by western blot (Fig. 5A). Although, phosphorylation of ATM, KAP1 and Chk1 was expectedly increased in HBEC-3KT cells treated with the DNA damaging agent etoposide, cells grown on TCP or 3 kPa did not show marks of an activated DDR at basal states [33]. Thus, a role for the DDR activation in the accumulation of HBEC-3KT cells in S phase was excluded.

**Fig. 5.**
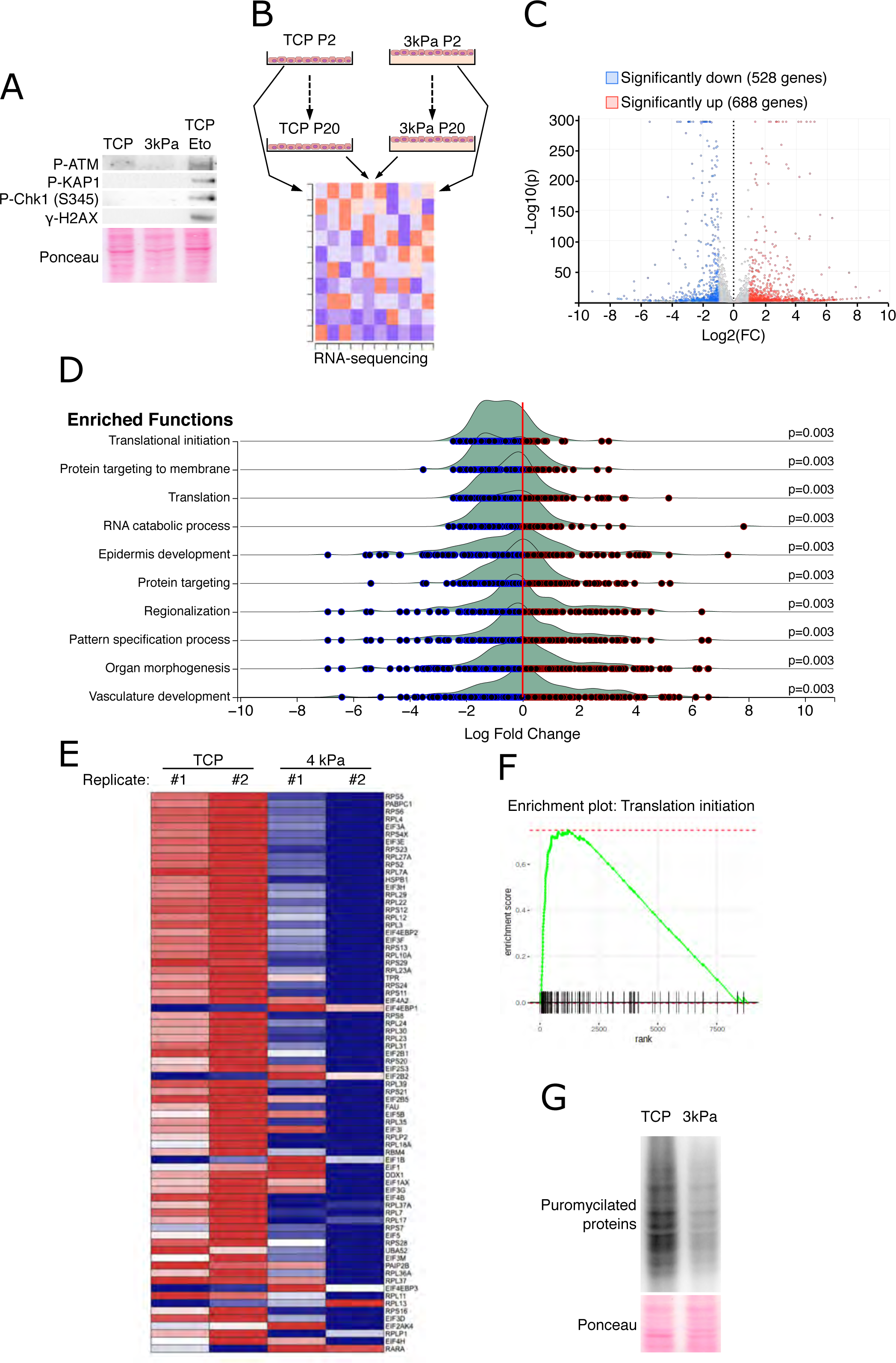
Pulmonary soft hydrogel does not trigger DNA damages but impairs translational processes and protein synthesis. A. Western blot analysis of markers of the DNA-damage repair (DDR) signaling pathways in HBEC-3KT cells cultured on TCP or 3kPa. Phosphorylated forms of ATM (P-ATM), KAP1 (P-KAP1) and H2AX (γ-H2AX) were used as markers the ATM pathway. Phosphorylated Ser345 of Chk1 (P-Chk1) was used as a reflect of the ATR axis. Cells grown on TCP and treated with etoposide (TCP Eto, 3 hours) were used as a positive control. Ponceau red staining was used as loading control. B. Schematic representation of the transcriptome analysis performed in HBEC-3KT cells cultured on either TCP or 3kPa supports. Technical details are provided in the material and method section. C. Volcano Plot of the differentially expressed genes (3kPa vs TCP, HBEC-3KT passage P2). Significantly downregulated genes are highlighted in blue while upregulated genes are in red. D. Ridgeline plots of the 10 most significantly enriched pathways. E. Heatmap Plot based on Translational Initiation gene signature for each condition. The pooled samples analyzed by individual sequencing are indicated by the replicate number. F. GSEA enrichment plot of the translation initiation gene signature. G. Protein synthesis measurement by SUnSET assay of HBEC-3KT cells cultured on TCP or 3kPa supports. Ponceau red Staining was used as loading control.

To obtain insight into the major pathways dysregulated on 3 kPa compared to TCP, RNA extracts from cells grown in both conditions were subjected to sequencing analysis at P2 and P20 (Fig. 5B). At P2, we observed a large number of genes whose expression is dysregulated by the soft matrix (Fig. 5C). Gene enrichment analysis unveiled a downregulation of factors associated to translation initiation pathway when the cells are grown on 3 kPa, suggesting an alteration of the protein synthesis process (Fig. 5D-F). Similar results were obtained at P20 (Supplementary fig. 2A, B). To functionally validate this translational impairment on soft substrate, a puromycin incorporation assay was performed on cells grown on both supports (Fig. 5G). Western blot analysis revealed a lower incorporation of this antibiotic in elongating peptide demonstrating a decreased protein synthesis rate in cells grown on 3 kPa supports.

### 2.5. Cells grown on soft supports exhibit an endoplasmic reticulum calcium leakage and activated ER stress at steady state

Since the RNA-sequencing data oriented towards a defect of the initiation step of translation, we next explored the underlying molecular pathway controlling the initiation of translation and potentially affected by the stiffness. The phosphorylation of eIF2a (p-eIF2a) is known to: i) limit the initiation step of translation and ii) favor a specific translational program, including the elevation of the amount of the transcription factor ATF4, both driving cell plasticity upon microenvironmental cues [34]. Furthermore, the phosphorylation of eIF2a is reported to trigger a cell cycle arrest in G1/S or G2/M upon chemical stressors [35,36]. We therefore investigated whether activation of this pathway could contribute to cell quiescence in S phase observed on soft supports. Western blotting analysis revealed an augmentation of p-eIF2a and ATF4 in cells grown on 3 kPa compared to TCP (Fig. 6A). These elevations are accompanied by an increase of BiP, a chaperone whose augmentation is a canonical marker of an activated ERS signaling. Furthermore, the expression of canonical ERS markers (*ASNS*, *TRIB3*, *CHOP* and *P58IPK)* was increased in cells grown on soft substrates compared to TCP (Fig. 6B). Furthermore, since ERS results from an impairment of protein homeostasis, the presence of aggresomes in cells grown on the soft supports was investigated (Fig. 6C). Ubiquitin-positive foci attested the occurrence of protein aggregates in these cells. Altogether, these results confirmed that cells grown on 3 kPa exhibit proteostasis defects and an active ERS at basal level.

**Fig. 6.**
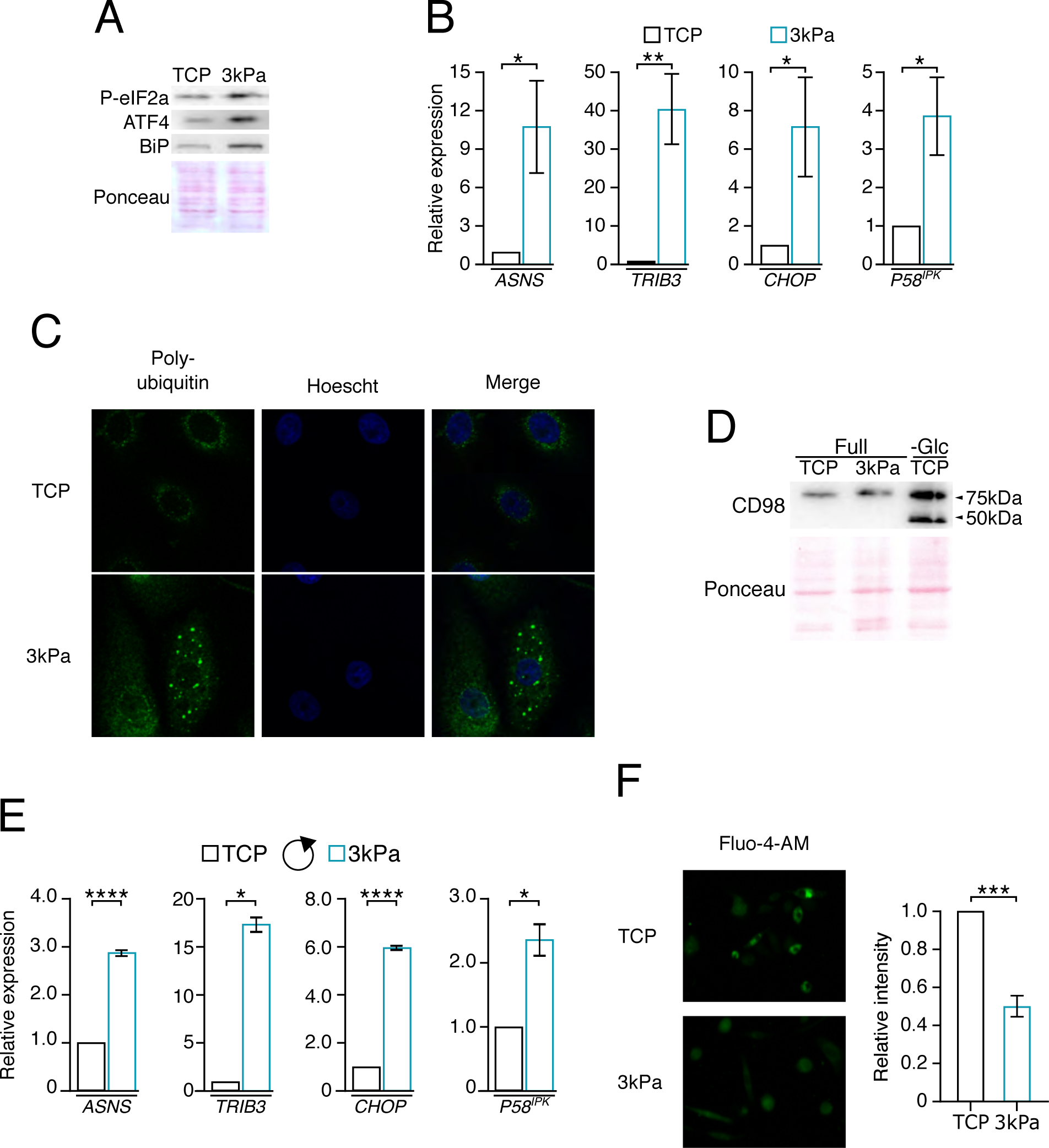
Pulmonary soft hydrogel rebalances Ca^2+^ concentration in the endoplasmic reticulum and triggers a basal ER stress. A. Western blot analysis of canonical ER stress markers (P-eIF2a, ATF4 and BiP) in HBEC-3KT cells cultured on TCP or 3kPa plates for 3 days. Ponceau red staining was used as loading control. B. Gene expression analysis of ER stress target genes *ASNS, TRIB3, CHOP* and *P58^IPK^* in cells seeded on TCP or 3kPa support. Data are expressed as mean +/-s.e.m of independent experiments (n = 4), Student’s t-test (*p< 0.05; **p<0.01). C. Analysis of aggresome formation in HBEC-3KT cells cultured on TCP or 3kPa supports. Immunofluores-cence staining against poly-ubiquitin (green) was performed after 3 days of cell culture. Hoechst dye was used to stain nuclei (blue). D. Western blot analysis of CD98 N-glycosylation in cells cultured on TCP or 3kPa plates for 3 days. HBEC-3KT grown on TCP and then deprived for glucose (-Glc, 24 hours) were used as a positive control. Pon-ceau red staining was used as a loading control. E. Gene expression analysis of ER stress markers (*ASNS, TRIB3, CHOP, P58^IPK^*) in HBEC-3KT cells grown on TCP or 3kPa for 3 days with daily medium refreshment (rounded arrow). Data are expressed as mean +/-s.e.m of independent experiments (n = 4), Student’s t-test (*p< 0.05; **p<0.01). F. Assessment of the intracellular calcium level in HBEC-3KT cells grown on TCP or 3kPa. After 3 days of culture, Ca^2+^ was stained using the Fluo-4AM dye (right panel). Relative fluorescence intensity was quan-tified (right panel). Data are expressed as mean +/-s.e.m of independent experiments (n = 3), Student’s t-test (***p< 0.001).

Then, we investigated the causality of the chronic ERS in cells grown on soft supports. Accumulation of protein aggregates classically results from either an impairment of the hexosamine biosynthetic pathway (HBP) and subsequent inhibition of protein N-glycosylation or a leakage of calcium (Ca^2+^) impairing the ER-chaperones folding activity [37,38]. We first tested whether a metabolic stress and subsequent glycosylation defects might be involved. To this end, the glycosylation of CD98, an adaptor of amino acid transporter and a model protein for assessing N-glycosylation process, was evaluated by Western blotting (Fig. 6D) [39]. Although glucose deprivation on TCP, impairing the HBP, consistently diminished the CD98 N-glycosylation as revealed by the appearance of a lower band, we did not observe such hypoglycosylated form of CD98 in cells grown on the 3 kPa supports, thus leading to the conclusion that the chronic ERS is not caused by a defect in glycosylation. Additionally, gene expression analysis of the ERS canonical target genes was performed after medium renewal in order to exclude any nutrient exhaustion that may have triggered the UPR (Fig. 6E). The expression degree of these markers remained higher in cells grown on soft substrates compared to TCP support Collectively, these results indicate that the ERS observed in cells grown on soft supports does not result from an alteration of the metabolic microenvironment. As mentioned above, the ERS can also be triggered through the impairment of the Ca^2+^ homeostasis within this organelle. Then we tested whether cells grown on 3 kPa supports display defects of Ca^2+^ storage within the ER. To this end, a chemical reporter was used for visualizing the free intracellular Ca^2+^. Validation experiment combining this chemical with an ER tracker confirmed that this ion is majorly localized in the ER lumen at basal state in the HBEC-3KT grown on TCP (Supplementary fig. 3). Then assessment of intracellular Ca^2+^ concentrations revealed a lower accumulation of Ca^2+^ in cells grown on 3 kPa compared to TCP providing a causal explanation to the chronic ERS observed on soft supports (Fig. 6F).

### 2.6. The eIF2a branch drives the S-phase quiescence on soft matrix and confers protection to ERS-mediated cell death by metabolic stress

Beyond its classical role of translation modulator, we then sought to establish whether the phosphorylation of eIF2a could be a cell cycle checkpoint controlling the S/G2 transition. To this end, cells grown on 3 kPa were treated with ISRIB, a compound blunting the stress response downstream the eIF2a phosphorylation [41,42][43]. ISRIB expectedly reduced the expression of ATF4 target genes (Fig. 7A). At the contrary, the mRNA level of P58IPK, whose expression is driven by the IRE1-XBP1 branch of the ERS, was not affected confirming ISRIB efficacy and selectivity [44]. Furthermore, the expression level of quiescence genes was also decreased to a degree closed to HBEC-3KT grown on TCP, indicating that p-eIF2a is contributing to the induction of quiescence markers in cells grown on soft supports. To determine the contribution of p-eIF2a in the cell cycle halt in S phase, cells grown on the soft substrates and treated or not with ISRIB and the consequences on the cell cycle were analyzed by flow cytometry (Fig. 7B). ISRIB promotes the cell cycle progression as it significantly lowers the accumulation of cells in the S-phase and we observed a trend of increased proportions of HBEC-3KT in G2/M and G0/G1 phases. These data demonstrate that alleviating the eIF2a phosphorylation permits the bypass of the S-phase quiescence in cells cultivated on soft supports.

**Fig. 7.**
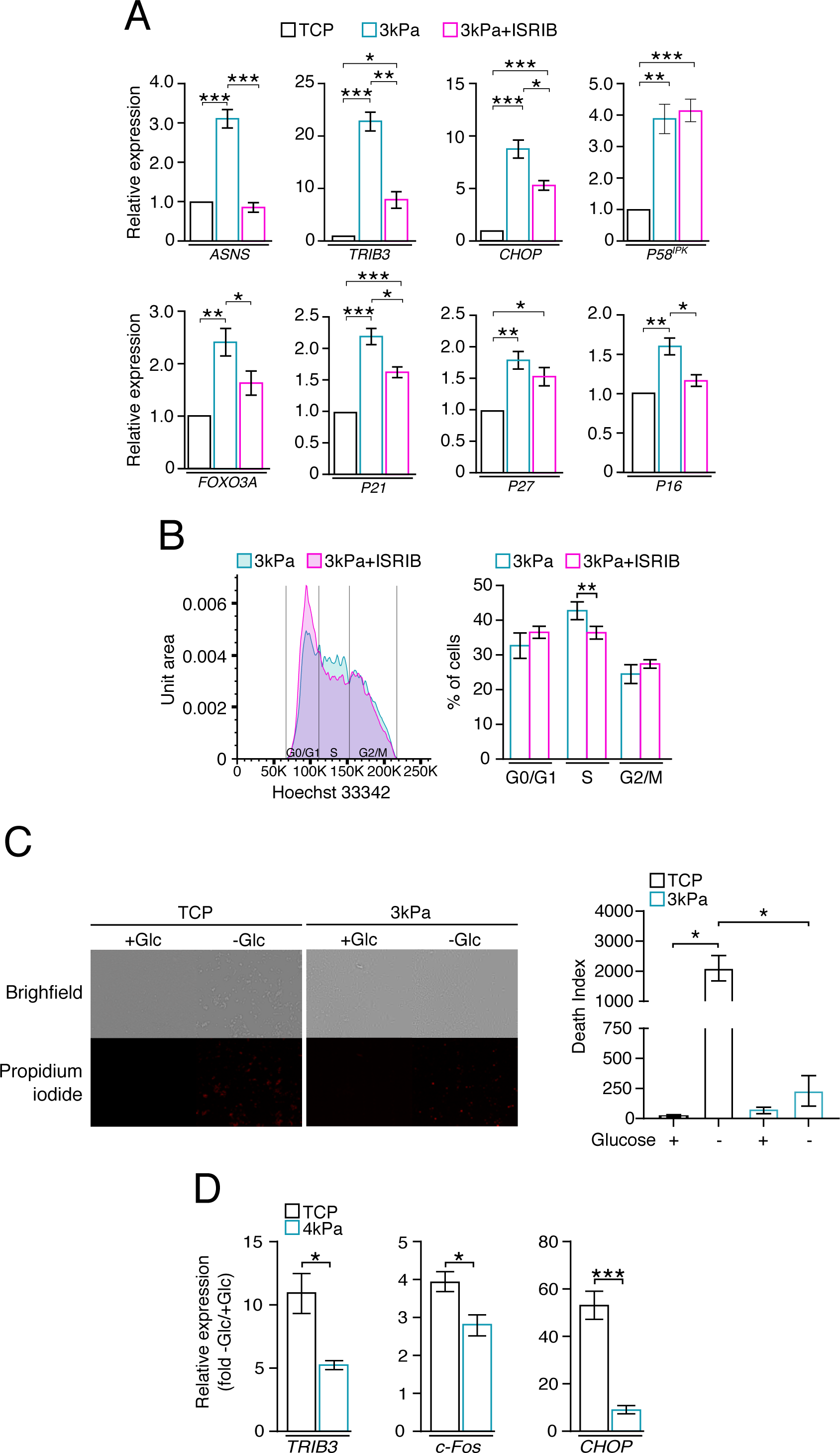
ISRIB treatment favors the S/G2 transition and soft matrix confers resistance to glucose deprivation. A. Gene expression analysis of *TRIB3, ASNS, CHOP, P58^IPK^, P16, P21, P27, FOXO3A* in HBEC-3KT cells platted in TCP or 3kPa and treated with or without ISRIB (400nM, 3 days). Data are expressed as mean +/-s.e.m of independent experiments (n = 8), One-way ANOVA and Tukey’s multiple comparisons test (*p< 0.05; **p<0.01, ***p<0.001). B. Cell cycle analysis by flow cytometry of cells cultured on 3kPa stiffness with or without ISRIB (400nM, 3 days). Cell distribution was plotted according to DNA content (left) and quantification of each cell cycle groups was assessed (right). Data are expressed as mean +/-s.e.m of independent experiments (n = 3), paired two-tailed t-test (**p< 0.01) C. Measurement of glucose starvation-induced cell death in HBEC-3KT cells grown on TCP or 3kPa stiffnesses. Following 3 days of culture, cells were subjected to media with or without glucose (25 vs 0mM) for 48 hours. After 48 hours, cells were imaged in brightfield for confluency and in red channel to assess propidium iodide fluorescence (dead cells) (left panel). Death index was quantified in each condition. Data are expressed as mean +/-s.e.m of independent experiments (n = 3), One-way ANOVA and Tukey’s multiple comparisons test (*p< 0.05). D. Gene expression analysis of ATF4-dependent proapoptotic genes (c*FOS, TRIB3, CHOP*) in HBEC-3KT cells grown on TCP or 3kPa and starved for glucose for 48 hours. Data are expressed as mean of fold change +/-s.e.m of independent experiments (n = 3), Student’s t-test (*p< 0.05; **p<0.01).

Finally, we intended to determine whether a chronic ERS could interfere with the cellular outcome upon microenvironmental cues and particularly glucose shortage. Lack of glucose is found in various diseases including cancer, ischemia-reperfusion or neurodegenerative diseases [45–47]. Upon irremediable hypoglycemic stress, ATF4 orchestrates the cell death by directly inducing the expression of proapoptotic genes including *CHOP, TRIB3* and *cFOS* [48,49]. However it has been shown that prior activation of the ERS and subsequent feedbacks involving P58^IPK^ inhibit the p-eIF2a signaling and prevent cells from the ATF4-mediated apoptosis upon ER stressors [50,51]. Thus, as a functional validation of the basal proteostasic stress occurring in cells grown on soft supports, we hypothesized that culture on pulmonary mechanomimetic supports might confer resistance to exogenous lethal ERS, including glucose starvation (Fig. 7C). To avoid any bias caused by differential glucose uptake and metabolism according to the tested rigidities, cells grown on 3 kPa or TCP were subjected to complete glucose starvation [52]. As expected, we measured an important cell death for cells grown TCP and starved for glucose. However this loss of viability was completely abrogated on soft supports demonstrating that this matrix and the subsequent molecular rewiring protects the cells from a lethal metabolically-induced ER stress. Consistently, transactivation by glucose deprivation of ATF4-proapoptotic target genes, including CHOP, was much lower in cells grown on 3 kPa compared to TCP (Fig. 7D). Overall these data demonstrate that the ERS signaling, particularly the eIF2a branch, drives the phenotypic changes of cells grown on soft matrix.

### 2.7. Mechanomimetic substrates maintain proteostasis dysfunction like that of human pulmonary basal cells

Overall, our data revealed that epithelial lung cells grown on 3 kPa polyacrylamide hydrogels coated with collagen I display profound molecular and biochemical reprogramming underlying cell state changes towards a basal identity. Defining whether these characteristics are artefacts or part of progenitor biology is crucial for validating the relevance of our findings in human and, in general the capacity of mechanomimetics supports to mimic the *in vivo* cell biology. Therefore, we first sought to identify the pathways specifically dysregulated in basal cells in the human pulmonary tissue. To this end, publicly available data from single cells experiments performed on human lungs were interrogated and gene ontology analyses were performed on the previously identified genes characterizing the different cell clusters of the airway (Fig. 8A) [53]. Our results confirmed that the major dysregulated processes in basal cells are closely related to translation and proteostasis reinforcing the translational characteristics of this cellular subpopulation. Then, we assessed whether an active ERS signaling is found specifically in these cells. To this end, we compared in the non-secretory contingent of epithelial cells the enrichment of the gene signature related to the UPR (Fig. 8B). Consistently, basal cells show a higher enrichment for the UPR signature compared to the differentiating basal and ciliated cells indicating that the UPR is a molecular determinant of the progenitor identity in the lung. Overall, these human lung tissue data confirm the relevance of our *in vitro* results and shows that the use of mechano-mimetic hydrogels as culture supports allows to closely reproduce the progenitor biology.

**Fig. 8.**
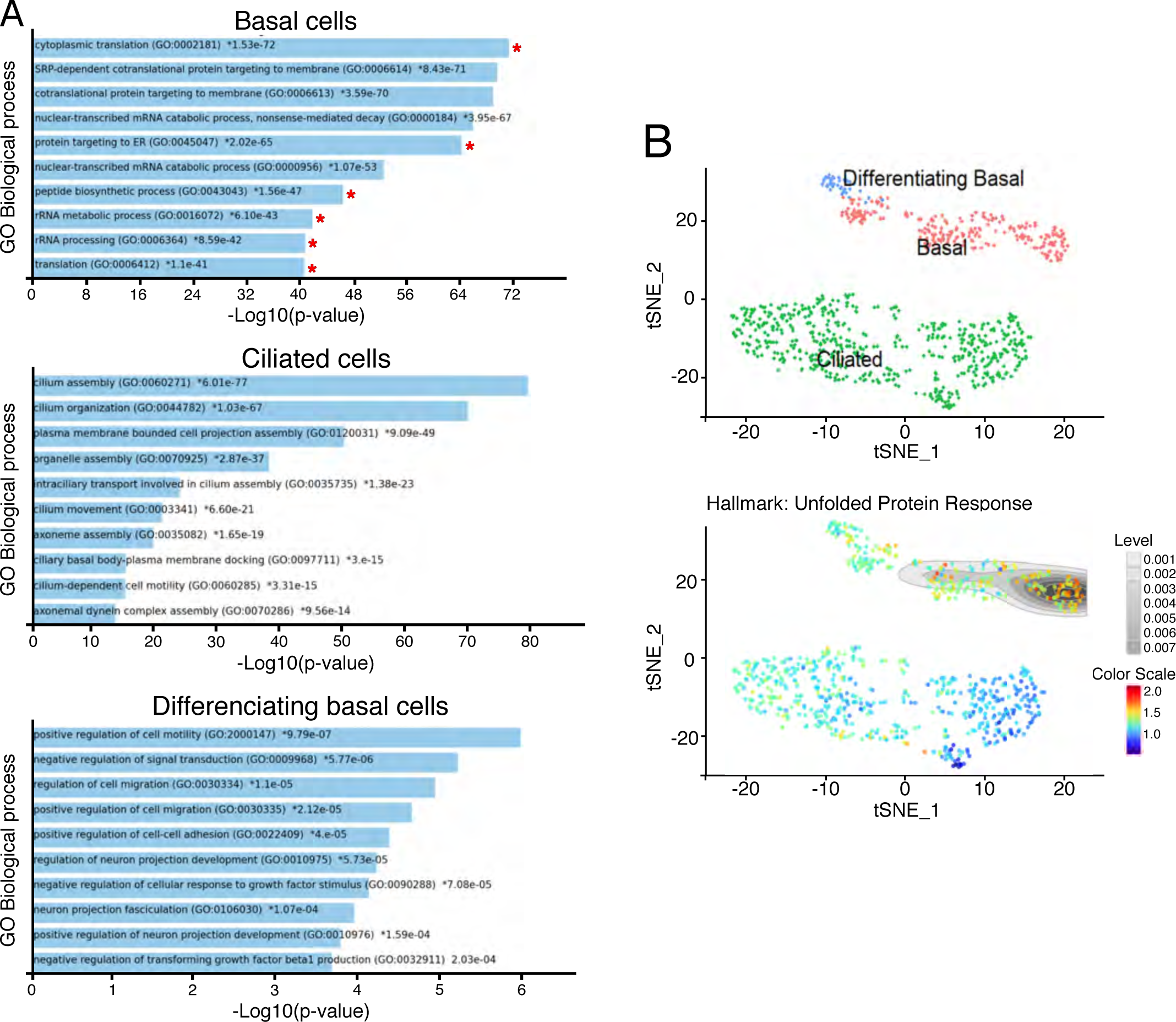
Pulmonary basal cells in human tissue exhibit marks of dysregulated translation and an active unfolded protein response. A. Bar chart of the top 10 enriched terms from the “GO_Biological_Process_2021” gene set library for basal, differentiating basal and ciliated cells. The enriched terms are displayed based on the −log10(p-value). The actual p-value is indicated next to each enriched term. B. t-SNE map depicting normal human basal cells (in red, 240 cells), differentiating basal cells (in blue, 43 cells), and ciliated cells (in green, 551 cells) (top panel). Visualization of the “unfolded protein response” signature (113 genes) in the t-SNE map of the 843 human cells represented above (bottom panel).

## 3. Discussion

The use of mechanomimetic supports for cell culture has gained an important interest in the last twenty years because of the development of mechanobiology and advances in understanding how the physical properties of tissues drive cell behavior or pathology development [11]. This technology has been consistently used to refine drug screening, disease modeling, and tissue regeneration. Nonetheless, technological issues regarding their fabrication and the need of a high degree of reproducibility, as well as the relevance of the cell biology towards the original tissue were still unclear.

Herein, we introduce an innovation oriented toward the design of soft-bottom multiwell plates to solve the issues associated with stiffness heterogeneity across wells while ensuring the impermeability of the wells. This development has facilitated the systematic examination of cellular proliferation over a span of 20 passages (about 7 months for the slowest condition) and allowed to conduct in parallel live imaging and biochemical and molecular analyses such as Western Blot, qPCR and RNA sequencing. The first observation was that HBECs were growing more than 3 times slower on supports with stiffness consistent with lung stiffness, of the order of 3 kPa (Figure 2). Although the impact of hydrogels on cell proliferation has been extensively reported [54], the type of quiescence induced by such supports remained unclear. In this study, we demonstrate that mechanomimetic supports trigger a drastic decrease of the S/G2 transition and an atypical quiescent state in S phase but not in the canonical G0/G1. This result differs from a previous study conducted on differentiated cells [55], where cells grown on soft matrices were shown to be halted in G1 phase. It also differs from the quiescence that is achieved by starving progenitor cells grown on TCP of serum[56] : in the latter case, the cells also showed a resistance to metabolic stress and the maintenance of their differentiation capabilities as we report, but with being halted in G1 phase. The unconventional quiescent state we report seems indeed to result from the combination of softness and progeny: it remains associated with the preservation of progenitor capabilities of basal cells and differentiation capabilities during passages. However, whether it is representative of the *in vivo* physiology remains a determinant for the validation of such supports in biology or as a relevant alternative to animal usage in preclinical studies. Incorporation of BrdU allows the detection of cells with active DNA replication and thus localizes in the S phase [57]. In lungs of mice and pork, the latter being proposed as a model closer to human tissue, the majority of pulmonary basal/stem cells are positive for BrdU whereas other epithelial cells show a very poor proportion of BrdU incorporation [58,59]. These data thus show that the positioning in the cell cycle of lung cells grown on the mechano-mimetic supports closely mimics the biology of progenitors *in vivo*, where these cells are positioned in the S phase. The S phase ensures DNA replication and the transition to the G2 phase. Maintenance of genome integrity is critical in dividing cells, especially progenitors, and consistently the S/G2 checkpoint is notably controlled by the ATR kinase sensing stressed replication forks [31]. Nonetheless, since we did not observe an elevation in molecular markers related to DNA damages and the ATR signaling pathway, we conclude that cell accumulation in the S phase does not result from alterations within the DNA. The latter is critical because amplification at a large scale of cell progenitors, including mesenchymal stem cells, without damaging the DNA is a key challenge in regenerative medicine and cellular therapy [60]. Instead of a contribution of the DDR pathways, we propose that soft substrates trigger the integrated stress response. We therefore investigated the phosphorylation of eIF2a as the latter actively contributes to the maintenance of the S-quiescence. The role of this molecular event in the cell cycle, particularly in progenitors, remains unclear. P-eIF2a has been proposed as a checkpoint for the G1/S or G2/M transition [35,36]. In these previous studies, the contribution of p-eIF2a to the cell cycle arrest was investigated using chemicals that might have pleiotropic effects and can artificially influence the phase in which the cell is arrested. Here we observe that cells grown on the 3 kPa substrates accumulate the phosphorylated form of eIF2a (Figure 6A). Thus, in contrast to previous studies, the underlying signal inducing p-eIF2a was generated by the non-genotoxic physiological mechanics, as attested by the absence of DDR activation. This indicates that the role of the eIF2a phosphorylation in the cell cycle arrest might depend on the context but is rather associated to a prolonged S phase in cycling cells upon physiological conditions. The relevance of the constitutively phosphorylated form of eIF2a in the normal physiology of cell progenitors is attested in several tissues. Indeed, p-eIF2a is abundant and promotes pluripotency of mouse and human embryonic stem cells [61]. Furthermore activation of the ER stress signaling is essential for the reprogramming of somatic cells into induced pluripotent stem cells (IPSCs) [62]. In line with these assumptions, p-eIF2a is implicated in the conservation of the stemness of hematopoietic cells and the maintenance of stem cell quiescence and self-renewal in the muscle of mice . This specific subpopulation of cells are not blocked by an arrest in G0/G1 but characterized by a prolonged S-phase of 14 hours, thus demonstrating that indeed, physiological ERS and p-eIF2a control the cell identity also *in vivo* [65].

Beyond the impact of the mechanics on cell fate, we herein unveil the calcium leakage from the ER and subsequent activation of the UPR signaling as a mechanical sensitive pathway. In accordance with a recent report by Chen et al. [66], our results demonstrate that physiological mechanics is sufficient to rebalance the intracellular calcium concentration but we additionally show that it triggers an ERS and subsequent molecular axes driving phenotypical changes. These data are consistent with the central role of the calcium flux in maintaining a quiescent state. Interestingly, modulation of the calcium efflux between the ER and the cytosol *in vivo* is sufficient to induce cell proliferation [67,68]. In the recent years, the ER has emerged as an integration hub for mechanical cues. Indeed, IRE1 and PERK, two key ER sensors, physically interact with the filamin A and F-actin [69,70]. These interactions remodel the cytoskeleton and dictate the cell migration properties. However how physical cues are integrated by the ER remains poorly described. Mechanical cues are sensed at focal adhesions which are integrin-based structures at the cell membrane. The ER-plasma membrane contact sites represent specific regions of interest for the integration of the physical cues, organelle contacts and calcium signaling. Interaction of STIM1, anchored in the ER membrane, and ORAI1, located within the plasma membrane, promotes calcium entry and limits the ER stress activity [71]. Assembly of the STIM1-ORAI1 complex is controlled by integrin signaling, notably the Wnt5a pathway which may be influenced by the mechanical microenvironment [72]. Further investigations are required to elucidate the molecular crosstalk between the physical microenvironment, regulation of calcium homeostasis and the cell identity regulation.

Our study also demonstrates that the mechanical environment protects against metabolic stress. Various pathologies combine profound rearrangements of the physical and nutritional microenvironment. Indeed, tissue stiffening is often associated with defects in angiogenesis and alteration of the tissue perfusion. How diseased cells survive to scarcity constitutes key challenges in the understanding of the mechanisms underlying pathologies development. This question must now be considered through the prism of the cellular identity since progenitors are classically described as: i) more resistant to cell death compared to differentiated epithelial cells and ii) the cell-of-origin of various chronic diseases such as cancer [73,74]. In line with our observation, multiple pathologies associate to physical changes of the tissue, also display an alteration of Ca^2+^ homeostasis and signaling associated to a decreased Ca^2+^ concentration within the ER lumen [75]. Additionally, antiapoptotic proteins such as BCL-2 reduces the concentration of Ca^2+^ in the ER providing resistance to apoptosis [76]. Consistently our results demonstrate that physiological stiffness and associated Ca^2+^ depletion is protecting against cell death caused by a lack of glucose and canonically attributed to a terminal ERS. This discrepancy of cell outcome upon stress depending on the stiffness of the extracellular environment cannot be attributed to a bias in glucose metabolism or nutrient exhaustion in the medium according to the substrate since cells were subjected to complete starvation at the same time. Thus, the protective effect observed on the soft substrate should be rather caused by mechanical-driven balancing of the Ca^2+^ concentration, priming progenitor cells to resist to a second ER stressors, limiting cell death of damaged cells and promoting disease development. So far, a large knowledge regarding stress response comes from mechanistical investigations performed on 2D culture grown on TCP, a rigid matter associated to aberrant proliferative rate and loss of cell identity. Revisiting these mechanisms on mechanomimetic supports, better reflecting the progenitor biology, would be a promising approach for the comprehension of cell outcome upon stresses and pathologies initiation and progression.

More generally, hits identification at early steps of drug discovery or toxicology assessments and mechanistical investigations imply the utilization of standardized and reliable cell culture methods compatible with prolonged exposure and high throughput screening and associated assays. 3D models do not currently comply with these requirements since spheroids/organoids: i) are costly to produce at large scale, ii) are often heterogenous in shape and size limiting their use by conventional imaging systems and iii) display a poor number of stem cells that exhausts along few passages leading to the loss of the culture [77]. This restricts their wide acceptance and use by the community and, thus, the traditional 2D culture still remains the gold standard with well-known limitations, notably in terms of cell drift. Nonetheless, although commercial culture media have been developed which balance metabolic fluxes as closely as possible to the physiopathology [78,79], the bias caused by the extraphysiological rigidity of plastic or glass remains poorly considered. Our results demonstrate the relevance of 2D mechanomimetic supports for preserving the molecular traits of progenitors and representing a practical tool for investigating the biology of this rare and hard-to-study subpopulation while preventing cellular drift for day-to-day cell-culture maintenance. Furthermore, our data argue in favor of the implementation of the mechanical microenvironment for evaluating stress response and cell outcome in more relevant conditions and improving the identification of compounds of interest.

## 4. Methods

### 4.1. Fabrication of mechanomimetics multiwells plates

Six-well plates with hydrogel bottoms were designed by Cell&Soft company (Grenoble, France). In brief, polyacrylamide hydrogels were photo-polymerized onto glass slides of dimensions 78 x 112 mm^2^ (Gataca Systems, Massy, France), following the protocol described in [26]. The duration of the UV exposure was chosen to achieve a stiffness close to 3 kPa. The gel was rinsed 3 times in deionized water to remove the unreacted compounds and left for swelling overnight. The gel was then dehydrated at room temperature. The plastic frame of a 6 wells bottom-less plate (Greiner Bio-One, Courtaboeuf, France, ref. 657160, home processed) was glued to the gel with a UV sensitive glue (NOA68, Norland Adhesives, Jamesburg, NJ, USA). NOA68 is based on polyurethane, a compound that is widely used in the medical industry. The surface of the gel was then rendered permissive to cell adhesion by coating Rat Tail collagen I (Gibco, Thermo Fisher Scientific, Whaltam, Massachusetts, USA, Ref. A10483-01) at a density of 1 µg/cm^2^ using the heterobifunctional crosslinker sulfo-LC-SDA (sulfosuccinimidyl 6-(4,40 -azipentanamido)hexanoate,Pierce, Thermo Fisher Scientific, Courtaboeuf, France) at a concentration of 0.6 mg/ml. The solution was removed after 1h and the gel was dehydrated for 45 min at room temperature. The gel was then exposed to UV light (UV KUB1, 16 mW/cm^2^, Kloe, Montpellier, France) for 5 min to bind the diazirine group of the crosslinker to the polyacrylamide gel. A solution of collagen 1 at 12 µg/ml was deposited on the surface and evaporated at room temperature under agitation, allowing the reaction of the NHS group of the sulfo-LC-SDA with the protein. This led to the theoretical crosslinking of 1 µg/ cm^2^ of Collagen I to the surface of the polyacrylamide hydrogel. The coated-soft plate could be stored dehydrated at 4°C up to three months. Culture treated plastic plates were coated with the same density of collagen I. Before use, the soft plate was extensively rinsed with PBS (Gibco, Thermo Fisher Scientific, Whaltam, Massachusetts, USA) and the gel was let for swelling 24 h at 4°C. PBS was replaced by culture medium 1 h before cell seeding, and the plates were placed at 37 °C in 5% CO_2_.The stiffness of the hydrogel was characterized before assembly with the bottom-less frame using indentation-type AFM as described in [26].

### 4.2. Cell culture, starvation experiments and reagents

The HBEC-3KT cell line was obtained from ATCC. This cell line was cultured in Keratinocyte Serum Free media (KSFM; Ref 17005-034, GIBCO, Thermo Fisher Scientific, Whaltam, Massachusetts, USA) supplemented with Bovine Pituitary Extract 50 mg.L^-1^ (Ref 13028-014, GIBCO, Thermo Fisher Scientific, Whaltam, Massachusetts, USA), glucose 16 mM (Ref A24940-01, GIBCO, Thermo Fisher Scientific, Whaltam, Massachusetts, USA), recombinant human Epithelial Growth Factor 5 µg.L^-1^ (Ref AF-100-15; PEPROTECH, Thermo Fisher Scientific, Whaltam, Massachusetts, USA) and Penicillin/streptomycin 1 % (Ref 15140-122; GIBCO, Thermo Fisher Scientific, Whaltam, Massachusetts, USA). If not specified, the experiments were performed at early passages P2–5).

Glucose starvation experiments were performed with Dulbecco’s Modified Eagle Medium devoid of glucose and sodium pyruvate (Ref A1443001, GIBCO, Thermo Fisher Scientific, Whaltam, Massachussets, USA) and supplemented with penicillin/streptomycin 1 % and 10 % dialyzed fetal bovine serum. The concentrations of nutrients were determined as previously described [80] : glucose 25 or 0 mM (ref A2494001, GIBCO, Thermo Fisher Scientific, Whaltam, Massachussets, USA) and with the subsequent amino acids purchased from Sigma Aldrich (Saint Louis, Missouri, USA) : L-proline 0.15 mM (ref P5607), L-alanine 0.15 mM (ref 05129), L-aspartic acid 0.15 mM (ref A8949), L-glutamic acid 0.03 mM (ref W328502), L-asparagine 0.34 mM (ref A4159) and L-glutamine 2 mM (ref G3126).

ISRIB (ref SML0843, Sigma Aldrich, Saint Louis, Missouri, USA) was bought in powder and dissolved in DMSO to be later used at a final concentration of 400 nM. Thapsigargin (Ref TQ0302, TARGETMOL, Wellesley Hills, Massachusetts, USA) was dissolved in DMSO and then used at 1 µM.

### 4.3. Air-Liquide Interface differentiation assay

Air-Liquide Interface cultures were performed according the recommendations from Stemcell technologies. In brief, cells were seeded at a density of 1.1×10^5^ cells/well onto a 12 mm Transwell (ref 38023, STEMCELL TECHNOLOGIES, Saint Égrève, France) and grown in KSFM until cell confluency. Then cells were subjected to air-lift. KSFM was removed from the upper and lower chambers and differentiation medium was added to the lower chamber only to allow contact with air. Differentiation medium was prepared as recommended by the company: PneumaCult-ALI Basal Medium was mixed with Pneumacult-ALI supplement (10X), Pneumacult-ALI Maintenance Supplement (100X), Heparin 4 µg/mL (Ref 07980, Stemcell technologies, Saint Égrève, France) and Hydrocortisone 480 ng/mL (Ref 07925, Stemcell technologies, Saint Égrève, France). Differentiation was monitored daily and differentiation medium was carefully changed every 2 days for 21 days. Differentiated ALI Culture of HBEC-3KT cells were then formalin-fixed, embedded in paraffin and processed for hematoxylin and eosin staining at the CRCL anatomopathological facility.

### 4..4. Cell cycle analysis

HBEC-3KT cells were plated (1 x 10^5^ cells/well) on TCP or 3kPa plates and grown for 3 days. Then cells were treated with ISRIB (400 nM) or vehicle for 3 days. Cells were trypsinized, washed with PBS and centrifuged at 300 g for 5 min. They were then counted and fixed in cold 70% ethanol for at least 2 hours on ice and pelleted. Cells were resuspended at 1 x 10^6^ cells/ml in staining solution, Hank’s buffered saline solution containing 4 µg/ml of Hoechst 33342 (Ref B2261, Sigma Aldrich, Saint Louis, Missouri, USA) for 1 h in the dark at 4°C. Stained cells were analyzed using a BD LSRFortessa™ flow Cytometer (BD Biosciences, Le Pont de Claix, France), and cell cycle distributions were determined with FlowJo software (Ashland, OR, USA).

### 4.5. Senescence-associated beta-galactosidase test

HBEC-3KT cells were plated (1 x 10^5^ cells/well) on TCP or 3 kPa plates and grown for 3 days. Cells were fixed in 0.5% glutaraldehyde solution (ref 49629, Sigma Aldrich, Saint Louis, Missouri, USA) for 10 minutes. beta-galactosidase solution was prepared containing 20 mM citric acid (pH=6) (ref 1.00244.0500, Merck Millipore, Darmstadt, Germany), 5 mM of potassium hexacyano-ferrate (II) trihydrate (ref 1.04894, Merck Millipore, Darmstadt, Germany), 5 mM of potassium hexacyano-ferrate (III) (ref1 .04973, Merck Millipore, Darmstadt, Germany), 2 mM of magnesium chloride (ref 1.05833, Merck) 150 mM of Sodium Chloride (ref S3014, Sigma Aldrich, Saint Louis, Missouri, USA) and 1 mg/ml X-Gal (ref EU0012-D, Euromedex, Souffelweyersheim, France) previously resuspended in N-N’ dimethylformamide. After fixation, cells were washed with beta-galactosidase solution to adjust cellular pH. Cells were incubated with beta gal solution and covered with aluminum for 6 hours at 37°C. Cells were then washed with PBS and nuclei was stained with Hoechst 33342 (Ref B2261, Sigma Aldrich, Saint Louis, Missouri, USA). Positive beta-galactosidase cells were then quantified and percentage was assessed using total nuclei number.

### 4.6. Intracellular calcium detection

HBEC-3KT cells were plated (1 x 10^5^ cells/well) on TCP or 3kPa plates and grown for 3 days. Fluo-4AM Solution (Ref FP-132303, Interchim, Montlucon, France) was prepared diluting Fluo-4AM (final concentration of 2µM) in 1:1 PBS-KSFM medium. Cells were then incubated with the mixture for 30 minutes and washed with Hank’s buffered saline solution for 30 minutes prior to imaging. The DPX blue/white ER-Tracker (ref 12353, Thermo Fisher Scientific, Whaltam, Massachussets, USA) was used to visualize the endoplamic reticulum in live cells. Images were acquired using Axio Vert.A1 microscope (ZEISS, Munich, Germany).

### 4.7. Cell Proliferation, cellular motility and cytotoxicity assay

For cell proliferation, HBEC-3KT cells were plated onto 6-well plate (1 x 10^5^ cells per well). Cell confluency was monitored and quantified over time using the Cellcyte X system (Cytena, Freiburg im Breisgau, Germany). Cells were also followed every 15 minutes over 2-hour long period to quantify cellular motility using the TrackMate Plugin for Fiji/ImageJ software (National Institutes of Health, Bethesda, Maryland, USA).

For cell death assessment, HBEC-3KT (1×10^5^cells/well) were seeded onto TCP or 3kPa 6-well plate and grown for 3 days. Then cells were subjected to indicated deprivation. Propidium iodide was added in the media at 2.5 µg/ml. Then, cell confluency and cell death were monitored over the indicated period of time using the Cellcyte X system (Cytena, Freiburg im Breisgau, Germany). Death index was calculated using the formula "(PI Foci Number/Confluency) x100”.

### 4.8. RNA extraction and RT-qPCR

Total cellular RNA was extracted after the indicated period of treatment using TRIzol Reagent (ref 15596026, Invitrogen, Carlsbad, California, USA) according to the manufacturer’s protocol. For cDNA synthesis, 0.25 µg of RNA were reverse transcribed using Superscript II reverse transcriptase (ref 18064014, Invitrogen, Carlsbad, California, USA) with random primers (S0142, Thermo Fisher Scientific, Whaltam, Massachussets, USA), according to the manufacturer’s instructions. cDNA was then amplified by qPCR using specific primers listed in supplementary Table 1 and the SYBR Green Master Mix (ref 1725274, Bio-Rad, Hercules, California, USA). qPCR was performed using the CFX connect real-time PCR system (Bio-Rad, Hercules, California, USA). Relative quantification was determined by using the delta-CT method. Expression of target genes was normalized against RPS11 mRNA levels used as an internal control.

### 4.9. Western blot analysis and SUnSET assays

To perform Western blot analysis, cells were platted on TCP or 3kPa plates (1 x 10^5^ cells/well) and grown for 3 days. Whole cell extracts were prepared from cultured cells lysed at 4°C in RIPA protein buffer containing protease and phosphatase inhibitors (ref 11697498001, Roche, Basel, Switzerland), and obtained by centrifugation at 13,000 g for 20 min at 4°C.

Protein concentrations of the cellular extracts were determined using the DC Protein Assay (ref 5000112, Bio-Rad, Hercules, California, USA). Equal amounts of proteins (20 μg) were separated by SDS-PAGE and then transferred onto nitrocellulose membranes (ref 1704271, Bio-Rad, Hercules, California, USA). Membranes were incubated in blocking buffer, 5% milk or Bovine Serum Albumin (BSA) in Tris-Buffered Saline/Tween 20 (TBST), for 1 h at room temperature, then incubated overnight at 4°C with the appropriate primary antibodies diluted in TBST containing 5% milk or BSA. Membranes were washed three times with TBST, incubated for 1 h at room temperature with the appropriate secondary antibodies, diluted in TBST containing 5% milk, and again washed three times with TBST. Detection by enhanced chemiluminescence was performed using the Immobilon Forte HRP Western substrate (ref WBLUF0500, Merck Millipore, Darmstadt, Germany) or SuperSignal West Femto Maximum Sensitivity Substrate (ref 34096, Thermo Scientific, Thermo Fisher Scientific, Whaltam, Massachusetts, USA). Red Ponceau was used as a loading control. The primary antibodies used were purchased from Bethyl laboratories: P-KAP1 (A300-2575A), from Cell Signaling Technology (Danvers, Massachusetts, USA): P-CHK1 (2348), Gamma H2AX (2577), CD98 (13180), BiP (3177), P-eIF2a (3398), ATF4 (11815), P-S6 (4858), from Merck Millipore (Darmstadt, Germany): P-ATM (05-740), Puromycin clone 12D10 (MABE343). The HRP-conjugated secondary antibodies (anti-rabbit and anti-mouse antibodies, respectively ref 7074 and 7076) were supplied by Cell Signaling Technologies (Danvers, Massachusetts, USA).

Nascent protein synthesis were evaluated using the surface sensing of translation (SUnSET) method as previously described by Schmidt et al. [81]. 15 min before being harvested and processed to prepare whole cell extracts in RIPA buffer, cells were incubated with 5 μg/mL of puromycin (ref P9620, Sigma Aldrich, Saint Louis, Missouri, USA) directly added into the medium. The amount of puromycin incorporated into nascent peptides was then evaluated by Western blot analysis on 20 μg of proteins using anti-puromycin antibody (ref MABE343) purchased from Merck Millipore (Darmstadt, Germany).

### 4.10. Immunofluorescence staining

The surface density of the collagen I was controlled by immunofluorescence using an Alexa Fluor 488 conjugated polyclonal antibody against the collagen I A1 (bs-10423R-A488, Cliniscience, Nanterre, France). Depth stacks obtained with confocal microscopy (Zeiss LSM880) were used as described in [82].

For aggresome assessment, HBEC-3KT cells were plated (1 x 10^5^ cells/well) on TCP or 3 kPa plates and grown for 3 days. Then cells were fixed in fixed in 4 % of paraformaldehyde in phosphate-buffered saline (PBS) before permeabilization with 0.5% Triton X-100 in PBS and stained with primary antibodies: anti-Ki67 (M7240, Agilent Dako, Santa Clara, CA, USA) diluted at 1/50 and anti-Poly-ubiquitin purchased from ENZO (ENZ-ABS840, Villeurbanne, France) diluted at 1/650. Secondary Antibody Alexa Fluor 488 Goat anti-mouse IgG (ref A-11029, Life Technologies, Carlsbad, California, USA) was used at 1:600. Nucleus were stained for 5 min with Hoechst n°33342 (ref H3570, Invitrogen, Carlsbad, California, USA) at 1:2000. Plates were mounted using the Fluoromount G mounting medium (ref 17984-25, EMS Diasum, Hatfield, Pennsylvania, USA). For Ki-67 staining, fluorescence microscope (Axio Vert.A1, ZEISS, Munich, Germany) was used to acquire image. For poly-Ubiquitin staining, ZEISS Confocal imager (40X) Zeiss LSM 880 was used. Final images were analyzed and cropped with the Fiji/ImageJ software (National Institutes of Health, Bethesda, Maryland, USA).

### 4.11. RNA sequencing

For RNA sequencing, the method of pooled RNA samples was used[83]. HBEC-3KT cells were cultured for 2 (P2) or 20 (P20) passages onto TCP or 3kPa plates and grown for 3 days. Six biological replicates were generated per condition. RNAs were extracted using NucleoSpin RNA kit (Macherey-Nagel, Düren, Germany). Then, for each condition, 3 replicates (1μg) were pooled thus providing 2 individual RNA samples treated individually until sequencing. Library preparation and RNA sequencing were performed at the ProfileXpert platform (Université Claude Bernard of Lyon, Lyon). Quality of samples was checked by Bioanalyzer 2100 (Agilent) and RNA was quantified by Quantifluor RNA kit (Promega, Charbonnières-les-bains, France). First, mRNA was enriched from 1 µg of total RNA, then library preparation was realized with the NextFlex Rapid Directional mRNA-Seq kit (Bio-Scientific, Perkin-Elmer, Villebon-sur-Yvett, France). Quality of libraries were checked by Fragment Analyzer (Agilent) and quantified by qPCR. Samples were put on Flow Cell High Output. Amplification and sequencing were performed with Illumina NextSeq500: run Single Read 76 bp was performed. After demultiplexing of the data with Bcl2fastq v2.17.1.14 (Illumina Inc, San Diego, CA, USA). Reads where mapped using Bowtie 2 on the human genome GRCh38 and counted using htseq count in the Galaxy project platform [84]. Differentially expressed genes were identified using DESeq2 and volcano plot were generated in ExpressAnalyst platform. All enrichment analysis were performed in ExpressAnalyst platform [85].

### 4.12. Single-cell data analysis

Already annotated, normal human single-cell RNA sequencing data set (“facs_normal_lung_blood_scanpy.20200205.RC4.h5ad”) were downloaded from https://www.synapse.org/#!Synapse:syn21560554 [53]. The scanpy object contained 9409 cells obtained during surgery from three patients undergoing lobectomy, including epithelial, endothelial, immune, and stromal cells. The sampled tissue corresponded to histologically normal lung tissue (bronchi, bronchiole, and alveolar regions), along with peripheral blood. Data were processed using R 4.2.0 and RStudio 2022.12.0+353. Downloaded scanpy object (.h5ad) was converted into a Seurat v4 object (.RDS) using “anndata” 0.7.5.6 and “Seurat” 4.3.0, prior to subsetting the cells of interest (“subset” function of Seurat)[86–88]. Cells of interest were represented by basal cells (three clusters, one from each patient), differentiating basal cells (one cluster), and ciliated cells (three clusters, one from each patient). We processed the new object using “Seurat” package, as follows: normalization (“NormalizeData” function), identification of highly variable features (“FindVariableFeatures” function), scaling the data (“ScaleData” function, performed on all the genes), linear dimensional reduction (“RunPCA” function), and non-linear dimensional reduction (“RunTSNE” function). All the clusters corresponding to the same cell type (for example, the three clusters of basal cells) were merged using the “RenameIdents” function. The t-distributed stochastic neighbor embedding (t-SNE) reduction was visualized using the “DimPlot” function. Differentially expressed features were calculated for each of the three cell types with the “FindAllMarkers” function, using the following parameters: only.pos = TRUE, min.pct = 0.25, logfc.threshold = 0.25, and kept only genes with an adjusted p-value <0.05. Then, we used “EnrichR” and “Enrichr Appyter” to identify and visualize top enriched terms [89,90]. Unfolded protein response gene signature was analyzed and plotted with “Single-Cell Signature Explorer” (https://sites.google.com/site/fredsoftwares/products/single-cell-signature-explorer). First, with the “Single-Cell Signature Scorer” software, we computed for each cell a signature score based on the “HALLMARK_UNFOLDED_PROTEIN_RESPONSE” signature from the “The Molecular Signatures Database”. Then, we used the “Single-Cell Signature Merger” software to collate the signature scores table with t-SNE coordinates, and, finally, the “Single-Cell Signature Viewer” software to display on a t-SNE map the signature scores .

### 4.13. Statistics

All of the statistical analyses were performed using GraphPad Prism version 6.0 (GraphPad Software, San Diego, CA, USA) by using Student’s t-test, Paired t-test, One-Way ANOVA or Two-way ANOVA and post-hoc Tukey multi-comparisons test. p-values less than 0.05 were considered statistically significant. Absence of mark “*” indicates lack of statistical significance.

## 5. Acknowledgements

We thank the D Bernard’s lab for technical assistance in senescence assessment and sharing reagents, B. Manship for English editing and Y. Chaix for funding management. The authors are grateful to the members of the Imagerie Cellulaire, cytometry and ProfileXpert facilities for their collaborative work. This work was supported by the ANR (ANR-22-CE51-0043-01) and INCa (PLBIO22-227), Cancéropôle CLARA (CVPPRCAN000174, CVPPRCAB000180 and CV-2021-039), Region Auvergne Rhone-Alpes (19-010898-01), Ligue Nationale contre le Cancer (R23031CC, R19007CC), Institut Convergence François Rabelais (17IA66ANR-PLASCAN-MEHLEN). A.N. acknowledges the support of the French Renatech network. A.N, C.M. and C.H. thank the microscopy facility MuLife of IRIG/DBSCI, funded by CEA Nanobio and GRAL LabEX (ANR-10-LABX-49-01) financed within the University Grenoble Alpes graduate school CBH-EUR-GS (ANR-17-EURE-0003).

## 8. Author contributions

PA L: Data curation, Methodology, Software., M P: Data curation, Methodology., P LG: Data curation, Methodology., M I: Data curation, Methodology, Validation., J F: Data curation, Methodology, Validation., I C: Supervision, Investigation., T R: Supervision, Investigation., N A: Validation Supervision, Investigation., S A: Validation, Supervision, Investigation., C H: Data curation, methodology. C M: Data curation, methodology, C D: Data curation, Methodology, Validation, Investigation., P B: Data curation, Methodology, Investigation., C FP: Validation Supervision, Investigation., A N: Data curation, methodology, conceptualization, funding acquisition, Writing-Original draft preparation., C C: Conceptualization, supervision, funding acquisition, Writing-Original draft preparation.

