## Supplementary data for "Soft extracellular matrix drives an endoplasmic reticulum stress-dependent S quiescence underlying molecular traits of pulmonary basal cells"

Supplementary information includes:

- Supplementary Figures 1 to 3
- Supplementary table 1

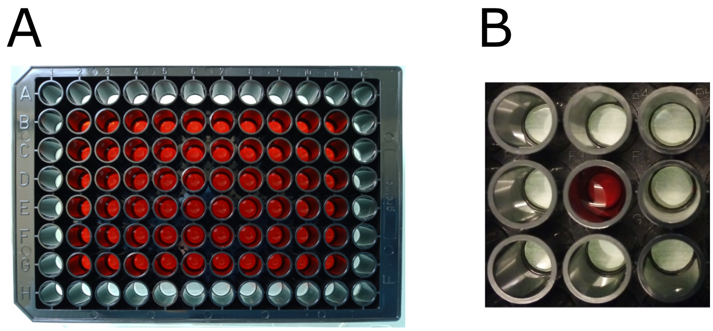

**Supplementary figure 1: 96-well soft supports fabrication according the presented method and visualization on absence of interwell leakage.**

A. The central wells of the assembled plate have been filled with colored water (Allura Red dye diluted in deionized water). The wells at the edge remain dry and clear.

B. Focus on a single well: the colored solution does not diffuse to the neighboring wells.

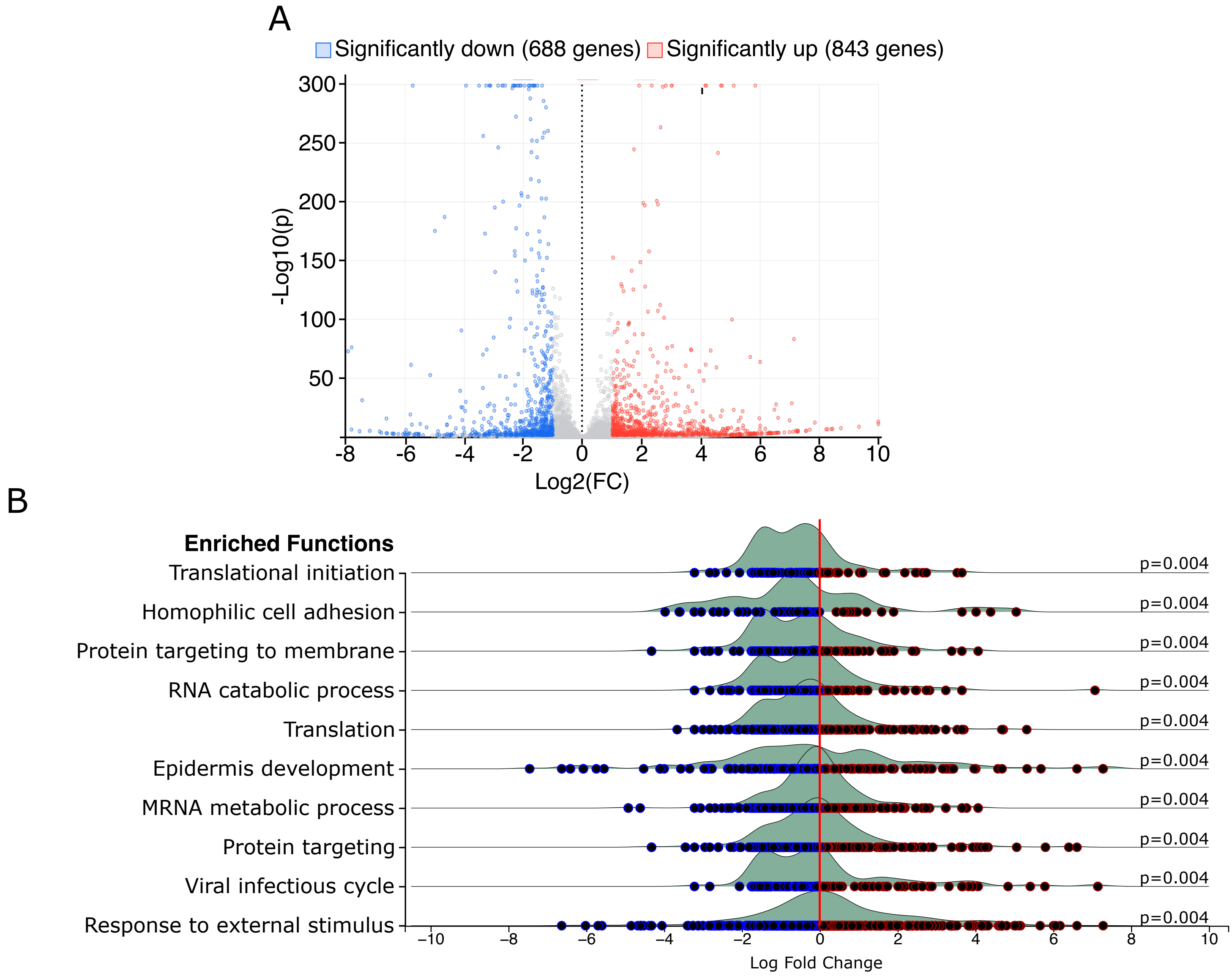


**Supplemental Figure 2: RNA seq data analysis of HBEC-3KT cells grown on TCP or 3kPa supports for 20 passages**

A. Volcano plot of the significantly down- and up-regulated genes in HBEC-3KT grown on soft vs TCP

B. Ridgeline representation of the GSEA analysis of the RNA seq data

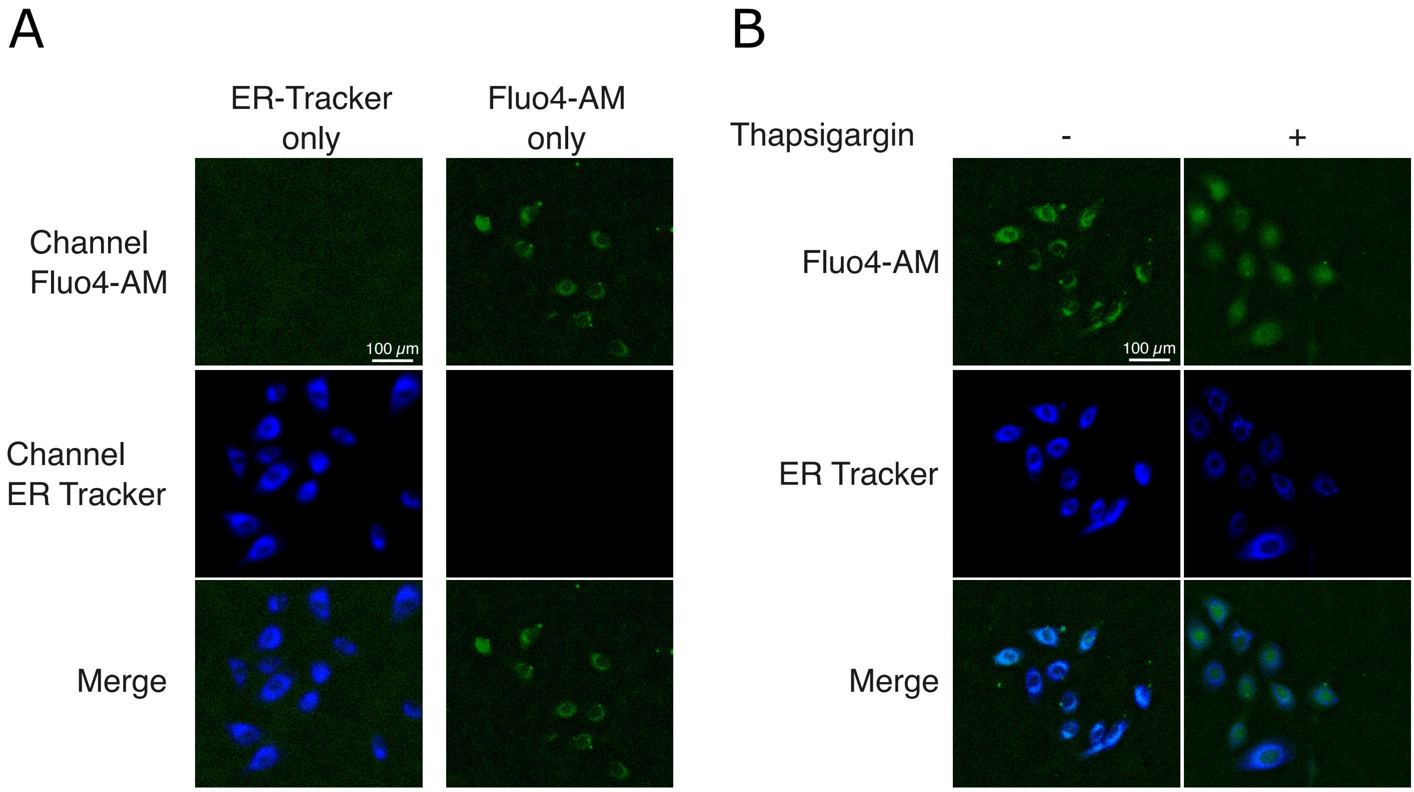


**Supplementary figure 3: Validation of the Fluo4-AM probe as a marker of the calcium in the endoplasmic reticulum.**

To assess whether the detected calcium is localized in the endoplasmic reticulum, HBEC-3KT cells were stained with the Fluo4-AM probe to stain intracellular calcium. The same cells were also stained with an ER tarcker to delineate the endoplasmic reticulum within the cell.

A. First, experimental parameters were determined by confirming that each visualized signal (ER tracker, Fluo4-AM) did not overlap with each other.

B. When combined, the chemical probes confirmed that free calcium in HBEC-3KT is restricted within the ER. Using the same parameters, cells were then exposed to the sarco/endoplasmic reticulum $Ca^{2+}$-ATPase (SERCA) inhibitor (thapsigargin 1μM, 30 min) and calcium diffusion from the ER was observed. t

| | |
|---|---|
| *P16* | Fw: CGGTCGGAGGCCGATCCAG<br>Rev: GCGCCGTGGAGCAGCAGCAGCT |
| *P21* | Fw: CACTGTCTTGTACCCTTGTGC<br>Rev: GGCGTTTGGAGTGGTAGAAAT |
| *P27* | Fw: TTTGACTTGCATGAAGAGAAGC<br>Rev: AGCTGTCTCTGAAAGGGACATT |
| *FOXO3A* | Fw: GATAAGGGCGACAGCAACAG<br>Rev: CGACTATGCAGTGACAGGTTG |
| *PDPN* | Fw: GTGCCGAAGATGATGTGGTGAC<br>Rev: GGACTGTGCTTTCTGAAGTTGGC |
| ACE2 | Fw: TCCATTGGTCTTCTGTCACCCG<br>Rev: AGACCATCCACCTCCACTTCTC |
| *KRT5* | Fw: GCTGCCTACATGAACAAGGTGG<br>Rev: ATGGAGAGGACCACTGAGGTGT |
| *KRT14* | Fw: TGCCGAGGAATGGTTCTTCACC<br>Rev: GCAGCTCAATCTCCAGGTTCTG |
| *TP63* | Fw: CAGGAAGACAGAGTGTGCTGGT<br>Rev: AATTGGACGGCGGTTCATCCCT |
| *ASNS* | Fw: CTGTGAAGAACAACCTCAGGATC<br>Rev: AACAGAGTGGCAGCAACCAAGC |
| *TRIB3* | Fw: TGGTACCCAGCTCCTCTACG<br>Rev: GACAAAGCGACACAGCTTGA |
| *CHOP* | Fw: GGTATGAGGACCTGCAAGAGGT<br>Rev: CTTGTGACCTCTGCTGGTTCTG |
| *P58IPK* | Fw: GCTGAATGTGGAGTAAATGCAG<br>Rev: GTGCAGCTTTTGATTTGCCC |
| *c-Fos* | Fw: CAGACTACGAGGCGTCATCC<br>Rev: TCTGCGGGTGAGTGGTAGTA |
| *RPS11* | Fw: AGCAGCCGACCATCTTTC<br>Rev: ATAGCCTCCTTGGGTGTCTTG |

**Supplementary table 1. List of human primers used for RT-qPCR experiments.**
